# Structural basis of broadly neutralizing human antibodies targeting diverse dengue virus forms

**DOI:** 10.64898/2026.02.07.704521

**Authors:** Ananya Chatterjee, Ankush Roy, Srividhya Srinivasan, Sylvia Charles, Jay Lubow, Leslie Goo, Vidya Mangala Prasad

## Abstract

Dengue virus (DENV) is a global health threat, with four antigenically distinct serotypes impeding vaccine and therapeutic development. We describe here, structural features of patient-derived, heavy chain–driven E-dimer–recognizing (HEDR) antibodies that achieve potent, pan-serotype activity by binding a conserved quaternary epitope on DENV’s envelope (E) glycoprotein dimer. Single-particle cryo–EM structures of three HEDR broadly neutralizing antibodies (bnAbs) complexed to DENV delineate epitope engagement via E-glycoprotein’s fusion loop region and conserved glycans. We also identify capsule-shaped DENV, with helical cryo-EM reconstructions of Fab-bound tubular DENV from different strains revealing that the E-glycoprotein arrangement is determined by virus strain, not bnAb identity. Our findings elucidate key binding features of HEDR bnAbs and signify the need for intervention strategies capable of neutralizing diverse DENV morphologies.

## Introduction

DENV is an enveloped RNA virus which affects nearly 400 million individuals worldwide giving rise to diverse clinical outcomes, including asymptomatic dengue fever, joint pain, rashes, dengue shock syndrome and severe dengue haemorrhagic fever(*1*). The canonical mature and infectious DENV is an icosahedral particle, 500 Å in diameter, with its outer membrane surface coated by its main antigenic protein, the envelope (E) glycoprotein (Fig.1A,B). Each E-protein monomer (∼50kDa molecular weight) consists of three domains—DI, DII, and DIII—with N-linked glycosylation at Asn67 and Asn153 amino acid residues(*2*, *3*). The E-protein’s DII domain contains the highly conserved fusion loop (∼14 amino acids long) at its distal end, which is crucial for viral membrane fusion within host cells during infection(*3–5*). On the mature virion surface, the E-proteins are arranged into 90 head-tail homodimers with three such E-dimers forming an E-raft that comprises two asymmetric units of the virus’s icosahedral surface arrangement (Fig.1B,C) (*2*). Despite global icosahedral symmetry, the local environment for each of the E-proteins in the asymmetric unit is different, and consequently, the local environment experienced by the E-proteins are distinct at the icosahedral 5-fold (5f), 3-fold (3f) and 2-fold (2f) symmetry positions (Fig.1B,C)(*2*).

**Figure 1.**
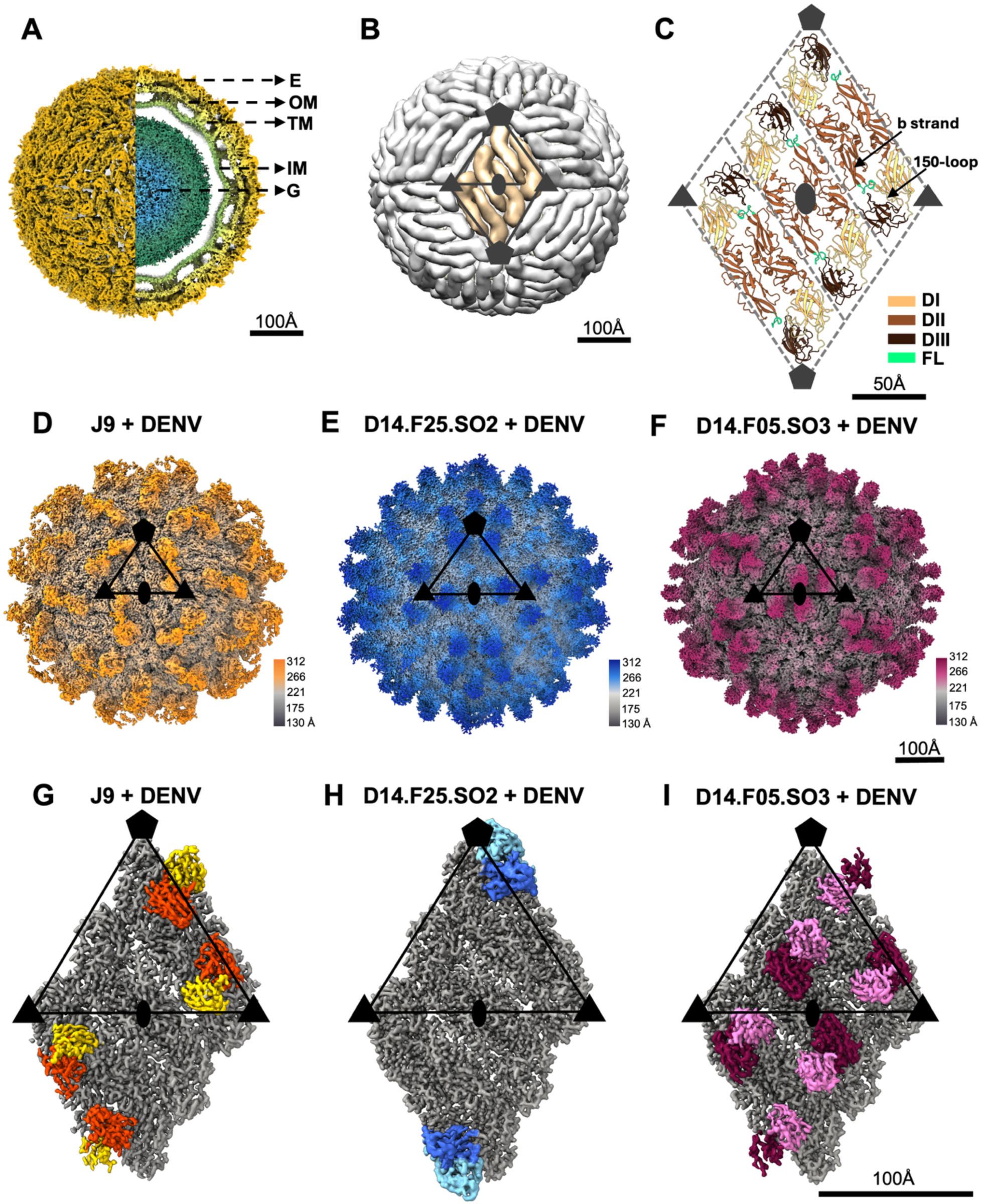
Cryo-EM structures of HEDR bnAbs complexed with DENV. **A**. Surface and cross-sectional view of dengue virus with black arrows indicating the different virus parts: E (envelope glycoprotein layer), OM (Outer lipid membrane), TM (Transmembrane space), IM (inner lipid membrane) and G (genome core). **B**. Surface representation of DENV virion showing icosahedral arrangement of E-proteins in grey. Black triangles indicate the icosahedral asymmetric unit. A single E- raft comprising of three E-dimers is coloured in beige and marked by a rhombus. **C**. Ribbon representation of an E-dimer raft, also referred to as an E-raft, consisting of three adjacent E-dimers. Parts of the E-protein are colored as: Domain-1 (DI) in mustard yellow, Domain-2 (DII) in brown with the fusion loop (FL) in green and Domain-3 (DIII) in black. **D-F**. Radially coloured, high-resolution cryo-EM density maps of DENV complexed with J9 (**D**), D14.F25.S02 (**E**) and D14.F05.S03 (**F**) Fab fragments. **G-I**. Density map representation of sub-particle reconstructions comprising the E-raft and Fab complexes of J9 (**G**), D14.F25.S02 (**H**) and D14.F05.S03 (**I**). In all panels, the 5-fold, 3-fold and 2-fold icosahedral symmetry vertices are indicated by a hexagon, triangle and ellipse, respectively.

DENV presents almost identical surface epitopes across its four serotypes (*6–8*), giving rise to cross-reactive antibodies that are, often, poorly-neutralizing against heterologous DENV strains(*9*, *10*). Secondary infections of DENV can, thus, cause severe disease due to antibody-dependent enhancement (ADE) of viral infection (*11*, *12*). There are few reported examples of broadly neutralizing antibodies (bnAbs) that neutralize all DENV serotypes (*8*, *13–15*). All these bnAbs bind on or near the conserved fusion loop region of DENV’s E-protein, highlighting the significance of this region as an immunogenic target. Notably, largest breadth in neutralization activity across DENV serotypes has been observed in antibodies that recognize a combination of the fusion loop along with adjacent regions on a E-protein dimer(*13–21*). From previously reported literature, few of the best characterized bnAbs which bind a quaternary epitope comprising the fusion loop and adjacent regions on the DENV E-protein are the E-dimer epitope (EDE) bnAbs, which are further sub-classified as EDE1 and EDE2 bnAbs(*19*, *20*). EDE1 bnAbs strongly neutralize both DENV and Zika viruses with bnAb binding being independent of N153 glycosylation on DENV E-protein. In contrast, the EDE2 bnAbs potently neutralize DENV, with lower activity against ZIKV, and require interaction with E-protein’s N153 glycan.

DENV populations in nature are a mixture of its mature and immature forms, including virions that are partially mature to various proportions(*6*, *22*, *23*). The mature and immature DENV structures are markedly different, with the mature DENV surface displaying 90 non-covalently bound, antiparallel E-dimers along with buried membrane (M) proteins on its surface, whereas the immature DENV displays 60 trimeric spikes of E-protein and precursor membrane (prM) protein heterodimers. Thus, incomplete virus maturation adds a layer of structural heterogeneity to circulating DENV populations. Additionally, in the past decade, DENV virions have been reported to deviate from their icosahedral structures to exhibit other morphologies such as the expanded/bumpy virus(*24*, *25*) and club-shaped tubular virions(*26*) at temperatures of 37° C and above. These heterogeneities in DENV structural forms exhibit variable proportions of distinct quaternary surface arrangements that are not yet fully understood, with little data available on the ability of bnAbs to recognize these various morphologies.

We describe here, high-resolution cryo-EM structures of three potent, broadly neutralizing human antibodies, J9, D14.F25.S02 and D14.F05.S03, in complex with the complete, infectious DENV virion. All three bnAbs were isolated from patients that recovered from secondary DENV infection(*14*, *15*). J9 and D14.F05.S03 neutralize only DENV whereas D14.F25.S02 neutralizes DENV as well as its relative, Zika virus (ZIKV). These selected bnAbs show strong neutralization activity against DENV which is comparable, and in some instances, better than other known bnAbs (*14*, *15*). The structures described here demonstrate that these bnAbs form a unique class of potent antibodies that bind to a quaternary E-dimer epitope encompassing the fusion loop region and conserved glycans, with distinct features that differentiate these from previously described bnAbs. We also report helical cryo-EM structures of tubular DENV morphologies bound to these bnAbs, showing the capability of these bnAbs to bind variant DENV morphologies. Furthermore, using different DENV strains, we demonstrate that E-protein arrangement on the helical DENV-bnAb structures is dictated by the virus strain and not the bound antibody.

### Cryo-EM structures of spherical DENV particles bound to bnAbs J9, D14.F25.S02 and D14.F05.S03

In this study, we have used two DENV-2 serotype strains from different geographical origins: THSTI/TRC/01(India) and US/BID-V594/2006 (Puerto Rico). Both these strains have been isolated from humans, and the respective E-glycoproteins share 98% identity (99% similarity) between each other and ∼98% identity (∼99.6% similarity) with the E-protein of well-studied DENV-2/16881 strain (Fig. S1). Using these strains, cryo-EM structures of spherical, mature DENV complexed with Fabs from three potent bnAbs, namely, J9, D14.F25.S02 and D14.F05.S03 were determined with icosahedral symmetry applied. The icosahedral cryo-EM structures of the DENV+J9 Fab, DENV+D14.F25.S02 Fab and DENV+D14.F05.S03 Fab were determined to highest global resolutions of 3.85 Å, 4.09 Å and 4.5 Å respectively (Fig.1D-F, Figs.S2-S7 and Table S1-S3). Further, sub-particle refinement and reconstruction of the E-dimer raft region improved the final resolution of the cryo-EM maps to 2.8 Å, 3.08 Å and 3.1 Å respectively for DENV+J9 Fab, DENV+ D14.F25.S02 Fab and DENV+D14.F05.S03 Fab complex structures (Fig.1G-I, Figs.S2-S7).

In the J9+DENV complex, sub-particle reconstruction of the E-raft showed clear Fab features near the icosahedral 5-fold (5f) and 3-fold (3f) symmetry positions, with no Fab density discernible at the 2-fold (2f) position (Fig.1G). In the D14.F25.S02+DENV complex, the overall Fab occupancy was very low, with distinct binding observed only at the 5f and 3f positions in the icosahedral reconstruction. Subsequent sub-particle reconstruction of the E-raft showed traceable Fab features only near the icosahedral 5f binding site, suggesting that the Fab binding was better at the 5f vertices (Fig.1E,H). In case of D14.F05.S03+DENV complex, sub-particle reconstruction of the E-raft showed clear Fab features near the icosahedral 3f and 2f binding sites, with fragmented Fab features near the 5f position, suggesting modestly lower occupancy of the Fab at the 5f binding position (Fig.1F,I). Thus, though one Fab (∼50kDa molecular weight) can bind to each E-protein for all three bnAbs, none of the bnAbs show full occupancy of all the possible 180 E-protein binding sites on the DENV virion, with each bnAb showing a different, preferential pattern of binding across the virion (Fig.1D-I) suggesting that complete occupancy of all available epitopes may not be necessary for neutralization activity of these bnAbs.

Despite variations in Fab occupancies, density for the Fab variable regions were well-resolved in each DENV+Fab complex at least at one binding site. The local resolution for the J9 and D14.F05.S03 Fab variable regions allowed clear interpretation of amino acid side chain densities. The local resolution for the D14.F25.S02 Fab variable region was marginally lower, likely due to the low occupancy of the fab, but still allowed confident tracing of the protein backbone and larger amino acid sidechains. The densities corresponding to the viral E and M glycoproteins were well resolved in all reconstructions, with clear amino acid side chains and glycosylation sites (N153 and N67) (Figs.S2,S4 and S6). These map features enabled real-space refinement of atomic models for the DENV E-protein and the associated Fab variable regions (Fig.1G-I, Figs.S2,S4 and S6, Tables S1-S3), permitting delineation of the antibody-antigen interface in all three Fab-DENV complex structures (Fig.2A-C).

**Figure 2.**
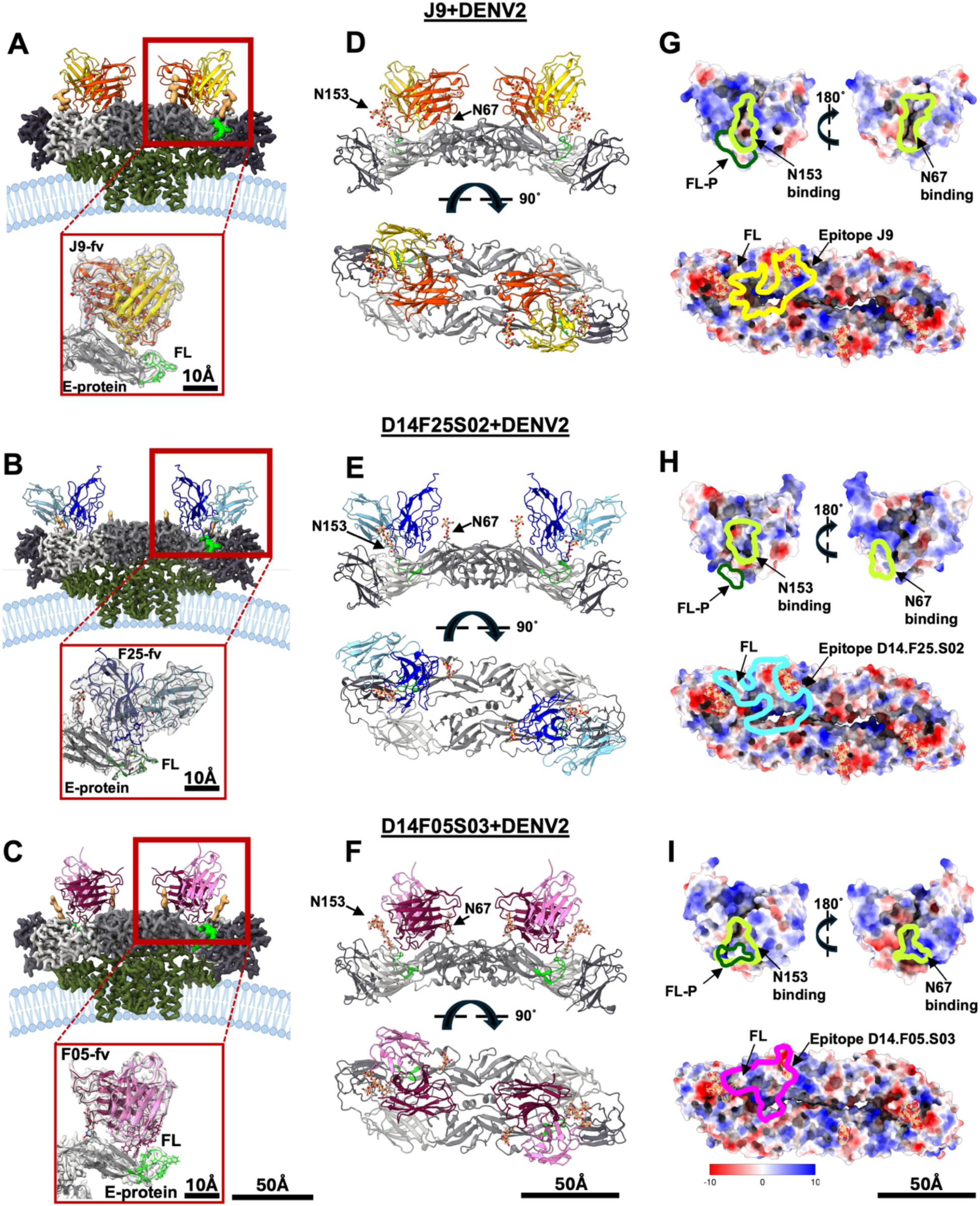
Quaternary binding epitopes of HEDR bnAbs on DENV E-dimer. **A-C**. Density map representation of DENV E-dimer (grey) and Fab complexes. Zoomed-in panel shows the built-in atomic model of the Fab in map density and its interaction with the E-protein fusion loop. The ribbon models of each Fab variable region (Fv) are coloured as: J9 Fv colored in orange (heavy chain) and yellow (light chain), D14.F25.S02 Fv colored in dark blue (heavy chain) and cyan (light chain), D14.F05.S03 Fv colored in dark pink (heavy chain) and light pink (light chain). E-protein is shown in grey and its fusion loop region in green. **D-F**. Top and side view of atomic models of E-dimer ectodomain complexed with Fab variable regions of J9 (**D**), D14.F25.S02 (**E**) and D14.F05.S03 (**F**) respectively. In all three atomic model representations, the glycan positions are marked as N153 and N67. **G-I**. Surface charge representation of the E-dimer epitope surface and the Fab paratope. Blue, white and red represent positive, neutral and negative surface charges respectively. The binding footprints are outlined for each Fab complex.

### Epitope features of bnAbs J9, D14.F25.S02 and D14.F05.S03 on spherical DENV

All three bnAbs directly interact with E-DII domain residues located in the b-strand (containing the N67 glycosylation site) and the fusion loop on one E-protein, along with residues in the 150 loop (containing the N153 glycosylation site) on the other E-protein within an E-dimer (Fig. 2A-F, Fig.S8). The bnAbs’ footprint, thus, span the E-dimer surface (Fig.2D-F), explaining why these were previously reported as not binding E-monomers in solution (*14*, *15*). The DENV-only binding bnAbs, J9 and D14.F05.S03, show the Fab positioned over the intra-E-dimer cleft with dominant interactions concentrated on and near the 150 loop region and the b-strand of adjacent E-proteins in an E-dimer (Fig.2, Fig.S8). In comparison, the DENV+ZIKV binding bnAb, D14.F25.S02, has the Fab positioned towards the interface between adjacent E-dimers, with dominant interactions seen on the b-strand and fusion loop, with few contacts to the 150 loop region (Fig.2, Fig.S8). Surface charge rendering of the bnAb paratopes and E-dimer surface shows a mixture of hydrophobic and electrostatic interactions influencing antibody binding for all three bnAbs (Fig. 2G-I).

Notably, all three bnAbs show overlapping epitope footprints, despite variations in their angle of approach to the E-protein and local differences in amino acid contacts (Fig.2, 3A, Fig.S8)(*14*, *15*). The overall epitope footprint of the three HEDR bnAbs (Fig.3A) also overlaps broadly with previously described EDE bnAbs(*20*), but the clusters of E-protein amino acid residues that are critical for bnAb binding and neutralization are distinct, as determined from these structural data and corroborating functional assays(*14*, *15*). In all the HEDR bnAb complex structures described here, the Fab heavy chain makes majority of interactions with the DENV E-dimer (Tables S4-S7) with the heavy chain of all three bnAbs involved in interactions with the fusion loop (Fig.2A-C, 3A). The HCDR3 of J9 and D14.F05.S03 is positioned near the fusion loop but angled towards the intra-E-dimer cleft, whereas the D14.F25.S02’s HCDR3 loop interacts in a sideway fashion with the E-protein’s fusion loop, such that the HCDR3 is inserted within the inter-dimer space between adjacent E-dimers (Fig.3A). Furthermore, all three bnAb heavy chain are also involved in interactions with conserved N153 glycan (Fig.3B) and the 150 loop on E-protein (Fig.2D-F, Fig.S8). This heavy chain-dominated recognition of epitope features groups these bnAbs as a separate class of their own, hereon referred to as heavy chain-driven E-dimer recognizing (HEDR) antibodies.

**Figure 3.**
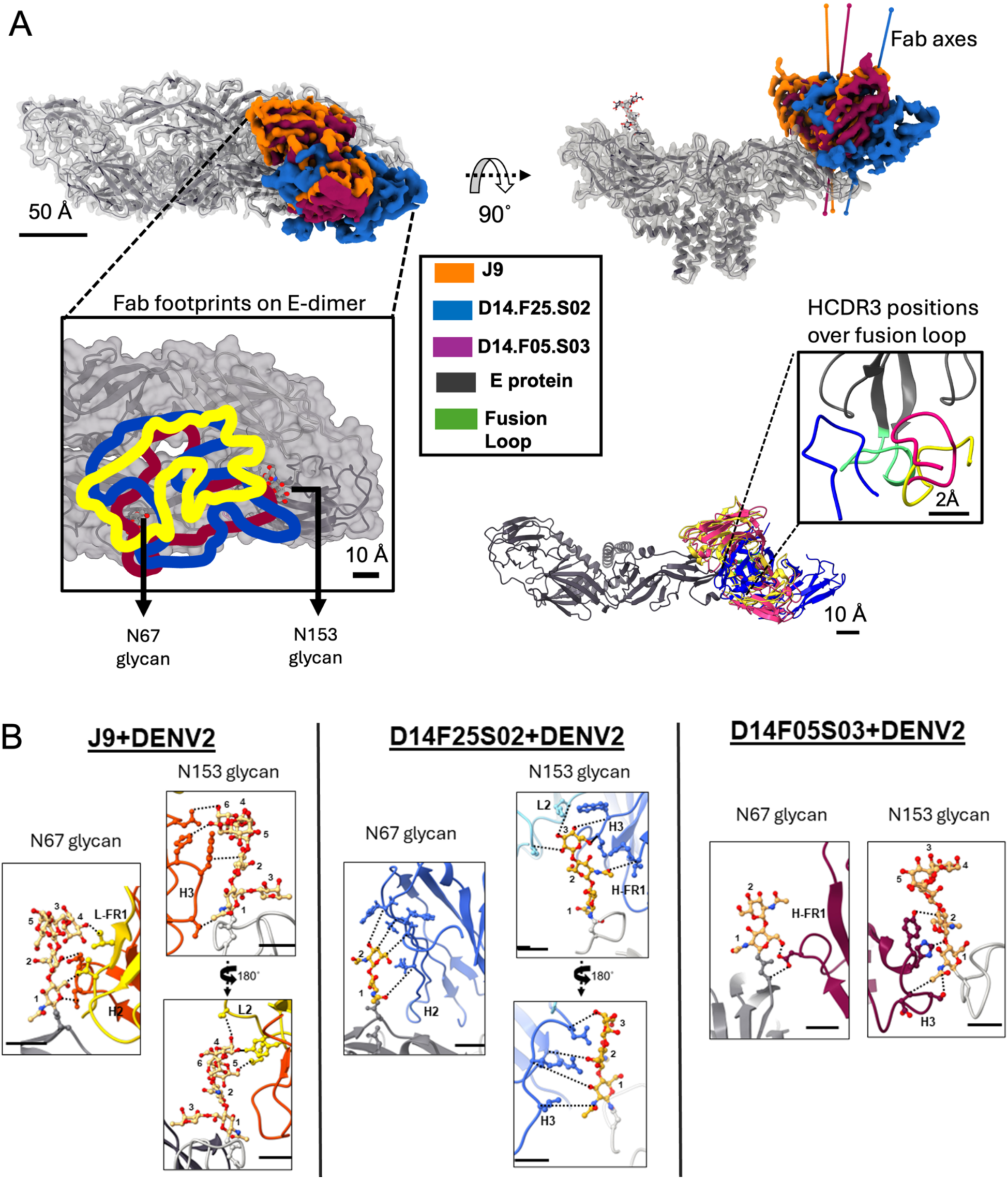
Overlap of HEDR bnAb binding epitopes and interaction with glycans. **A**. Top panels represent the top and side view of map density volumes of all three Fab variable regions (J9, D14.F25.S02 and D14.F05.S03) on E-dimer showing their overlapping binding positions. Zoomed-in panel in left bottom presents the top view of bnAb footprints on one of the E-dimer binding sites, showing the nestling of the bnAb footprint between the two conserved glycan positions. The bnAb footprint outlines are colored as J9: yellow, D14.F25.S02: blue and D14.F05.S03: magenta. Right bottom panel shows the binding position of Fab variable regions on the E-protein monomer, rendered as a ribbon diagram. Zoom-in panel represents a top view showing the positioning of the HCDR3 loops from all three bnAbs with respect to the fusion loop. **B**. Interactions of bnAbs J9 (left), D14.F25.S02 (middle) and D14.F05.S03 (right) with conserved glycans at N67 and N153 on E-protein. Black dotted lines indicate the closest interacting residues. Details on the interacting residues are given in Table S4-S6. Scale bar indicates 5 Å.

### Interactions with E-protein glycans by the HEDR bnAbs, J9, D14.F25.S02 and D14.F05.S03

The HEDR bnAbs can be classified into two categories, similar to the EDE bnAbs(*19*, *20*), based on their dependence on E-protein’s N153 glycan for neutralization activity. D14.F25.S02 forms HEDR category-1 which is independent of N153 glycosylation for its neutralization activity. Nevertheless, D14.F25.S02 bnAb makes multiple contacts with the N153 glycan (Fig3B, Table S5), in contrast to the previously described EDE1 bnAbs which induce disordering of the N150 loop containing the N153 glycan (*17*, *20*). D14.F25.S02 also uses its heavy chain to interact with the E-protein’s N67 glycan, which is also not essential for the bnAb’s neutralization activity (Fig. 3B). Thus, despite not requiring glycans for its activity, D14.F25.S02 accommodates both conserved glycans for binding to its epitope. On the other hand, HEDR category-2 bnAbs include J9 and D14.F05.S03, show many similarities with the EDE2 bnAbs, including the requirement of N153 glycosylation for its neutralization activity(*14*, *19*). J9 and D14.F05.S03 make close contacts to the N153 glycan making it an integral part of the bnAb epitope (Fig.3B, Table S4 and S6). J9 uses its HCDR3 to make predominant contacts with the N153 glycan, but the light chain also makes contacts with the terminal glycan residues (Fig.3B, Table S4). In case of D14.F05.S03, only the Fab HCDR interacts with the N153 glycan (Fig.3B, Table S6). J9 and D14.F05.S03 also make close contacts with the E-protein’s other conserved glycan at N67 position (Fig 3B, Table S4 and S6), though this glycan is not critical for its neutralization activity. As the spatial position of N153 glycan in DENV’s E-protein is different with respect to the equivalent N154 glycan position in ZIKV’s E-protein, the structural data suggests that close interactions with N153 glycan may influence differential recognition of DENV by the HEDR category-2 bnAbs, J9 and D14.F05.S03.

### Morphological features of tubular and capsule-shaped DENV present in virus preparations

Virus preparations used in this study were produced by infection of mosquito cells and incubated with Fabs at room temperature without any treatment at higher temperatures. Nonetheless, DENV particles with tube-like morphologies and bound Fab were observed as a small percentage (<15%) in all the cryo-EM datasets collected for both the US/BID-V594/2006 as well as THSTI/TRC/01 strains. Individual virions were captured from both DENV strains with different lengths of tube-like extensions connected to a spherical head (Fig.S9, S11), reminiscent of the club-shaped DENV morphology described previously(*26*). Furthermore, in case of the DENV2-US/BID-V594/2006 strain, completely tubular virions similar to cylindrical capsules were also observed (Fig.4A,B). The capsule-shaped DENV particles have similar widths as the tube-like extensions observed developing from spherical virions of the same strain, suggesting that capsule-shaped virions form via alterations to the spherical virus (Fig.S9). The capsule-shaped virions have an inner diameter of 110 Å and 120 Å in the D14.F25.S02- and J9-bound datasets respectively, with average length of particles being 79nm and 74nm respectively (Fig.4A,B). Thus, the capsule-shaped virions enclose a volume large enough to contain 1.9 – 2.2 copies of the viral genome (see Methods for details), indicating that these morphological variants can accommodate the viral genome in its entirety.

**Figure 4.**
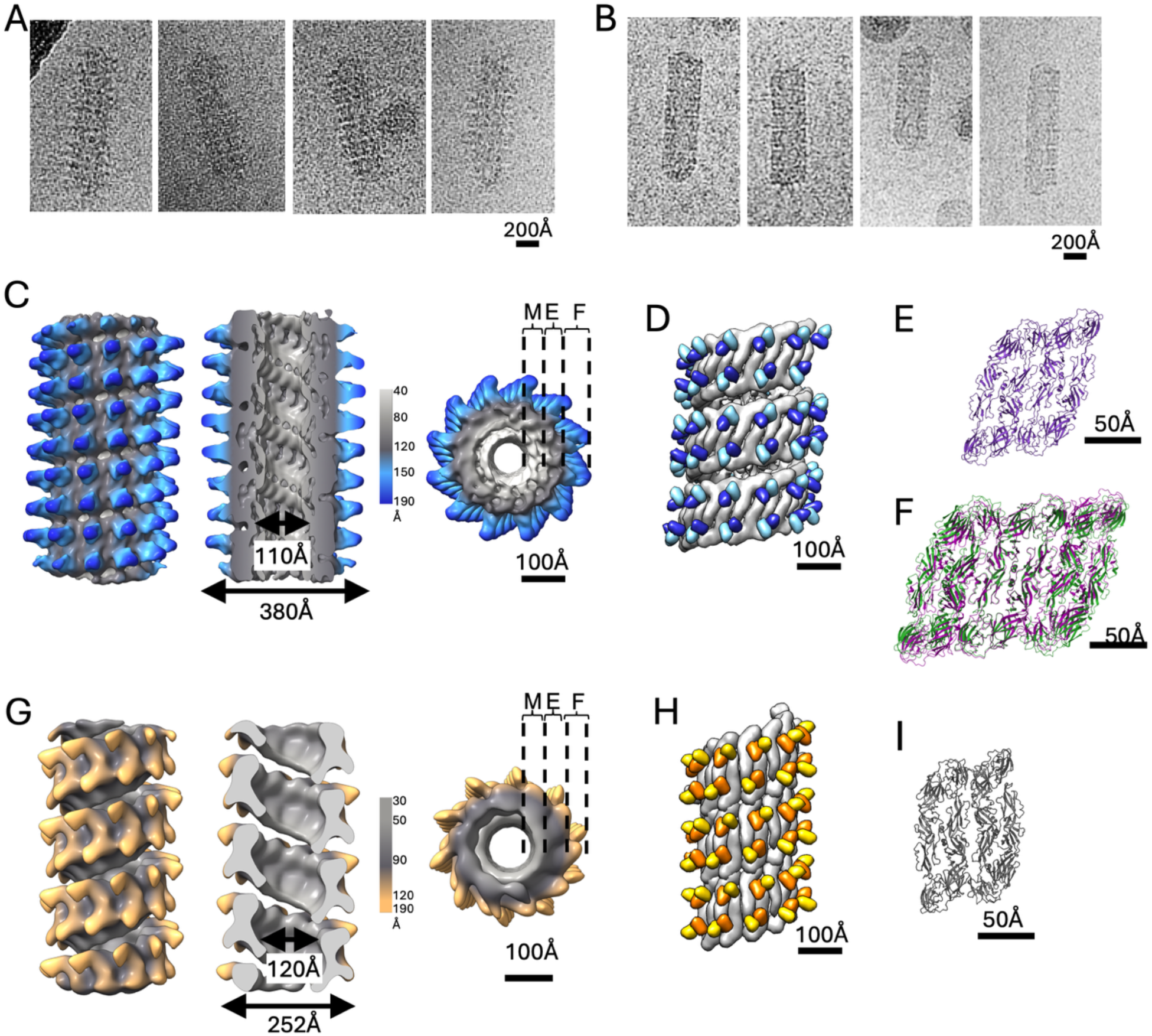
Capsule-shaped DENV show helical symmetry arrangement in Fab-bound state. **A-B**. Representative raw cryo-EM images of capsule-shaped DENV2-US/BID-V594/2006 virions bound with D14.F25-S02 Fab (**A**) and J9 Fab (**B**). Spiky Fab projections perpendicular to the smooth virus surface is observed along with presence of ordered protein arrangement. **C**. Map representation of surface (left) and longitudinal cut-out (centre) and top view (right) of the helical reconstruction of D14.F25.S02 Fab complexed with DENV2-US/BID-V594/2006. M, E and F represent the membrane, E-protein and Fab density regions, respectively, in the density map. **D**. Surface representation of fitted models of the E-dimer and D14.F25.S02 Fab variable region into the Fab-bound DENV2-US/BID-V594/2006 helical structure. **E**. Ribbon diagram of adjacent E-dimer positions in the D14.F25.S02 + DENV2-US/BID-V594/2006 helical structure. **F**. Overlap of three adjacent E-dimer atomic models from the D14.F25.S02 + DENV2-US/BID-V594/2006 helical structure (purple) to an E-raft from the DENV2-US/BID-V594/2006 icosahedral structure (green), showing similarity in E-dimer arrangement with an r.m.s.d value of 1.04Å. **G**. Map representation of surface (left) and longitudinal cut-out (centre) and top view (right) of the helical reconstruction of J9 Fab complexed with DENV2-US/BID-V594/2006. M, E and F represent the membrane, E-protein and Fab density regions, respectively, in the density map. **H**. Surface representation of fitted of the E-dimer and J9 Fab variable region into the Fab-bound DENV2-US/BID-V594/2006 helical structure. **I**. Ribbon diagram of adjacent E-dimer positions in the J9 + DENV2-US/BID-V594/2006 helical structure. This E-dimer arrangement is similar to that seen in the D14.F25.S02+DENV2-US/BID-V594/2006 helical structure (E) after rotation of 26° along an axis perpendicular to the plane of the E-dimer. R.m.s.d between the two E-dimer arrangements in panels E and I (after superposition with rotation) is 0.875Å.

### Cryo-EM structures of tubular DENV2-US/BID-V594/2006 virions bound to D14.F25.S02 and J9 Fab fragments

Cryo-EM reconstruction of the tubular regions of DENV2-US/BID-V594/2006+D14.F25.S02 Fab complex using helical symmetry parameters achieved a global resolution of 14.7 Å (Fig.4C; Fig.S9A-F, S10A). In this structure, E-dimers are arranged one next to another with a gentle rise, wrapping tightly around the membrane tube (Fig.4C-E, Fig.S9 A-E), with two Fabs bound per E-dimer, in similar positions as in the icosahedral structure. Overlapping three adjacent E-dimer atomic models from the helical structure with the icosahedral E-raft structure shows good agreement with an r.m.s.d value of 1.04 Å, confirming that the E-raft contacts seen in spherical virions are preserved in the helical structure of DENV2-US/BID-V594/2006+D14.F25.S02 Fab complex (Fig.4E,F). This helical arrangement of E-proteins also resembles the E-protein arrangement previously reported in the helical structure of ZIKV, a close relative of DENV, complexed with an EDE1 antibody(*26*). Similar helical patterns were observed in the J9+US/BID-V594/2006 dataset also, but only a very low resolution helical cryo-EM reconstruction was determined due to few tubular particle numbers (Fig.4G; Fig.S9 G-I, S10B). Rigid body fitting of the E-dimer and Fab complex model into the J9+DENV2-US/BID-V594/2006 map (Fig.4H, Fig.S9I) showed that the E-dimer arrangement is similar to that observed with the D14.F25.S02+DENV2-US/BID-V594/2006 helical reconstruction, albeit with a rotation of 26° along an axis perpendicular to the plane of the E-dimer (Fig.4E).

### Cryo-EM structures of tubular DENV2-THSTI/TRC/01 virions bound to J9, D14.F05.S03 and D14.F25.S02 Fab fragments

In the DENV2-THSTI/TRC/01 strain sample, only Fab-bound particles with a spherical head and narrow tube-like extension were observed (Fig.S11B,F, S12A). The tubular extensions were selected and subjected to unsupervised 2D classification, which also showed clear helical patterns (Fig.S11B,G 12B). Helical cryo-EM reconstructions of THSTI/TRC/01 virions were achieved to global resolutions of 14.41Å for J9-bound, 17.3Å for D14.F05.S03-bound and 17.37 Å for the D14.F25.S02-bound particles respectively (Fig.5, Fig.S11-S13). The outer and inner diameters of THSTI/TRC/01 tubular particles were 312 Å and 20 Å for the J9-virus complex, 280 Å and 55 Å for D14.F05.S03-virus complex, and 320 Å and 20 Å for the D14.F25.S02-virus complex respectively (Fig.5). The narrow inner diameters of the tubes in the THSTI/TRC/01 strains cannot accommodate the viral genome, indicating that the viral RNA is contained in the spherical head regions only. The narrow inner tube diameter could, thus, restrict these virions from transitioning to complete cylinders or capsule-shaped particles.

**Figure 5:**
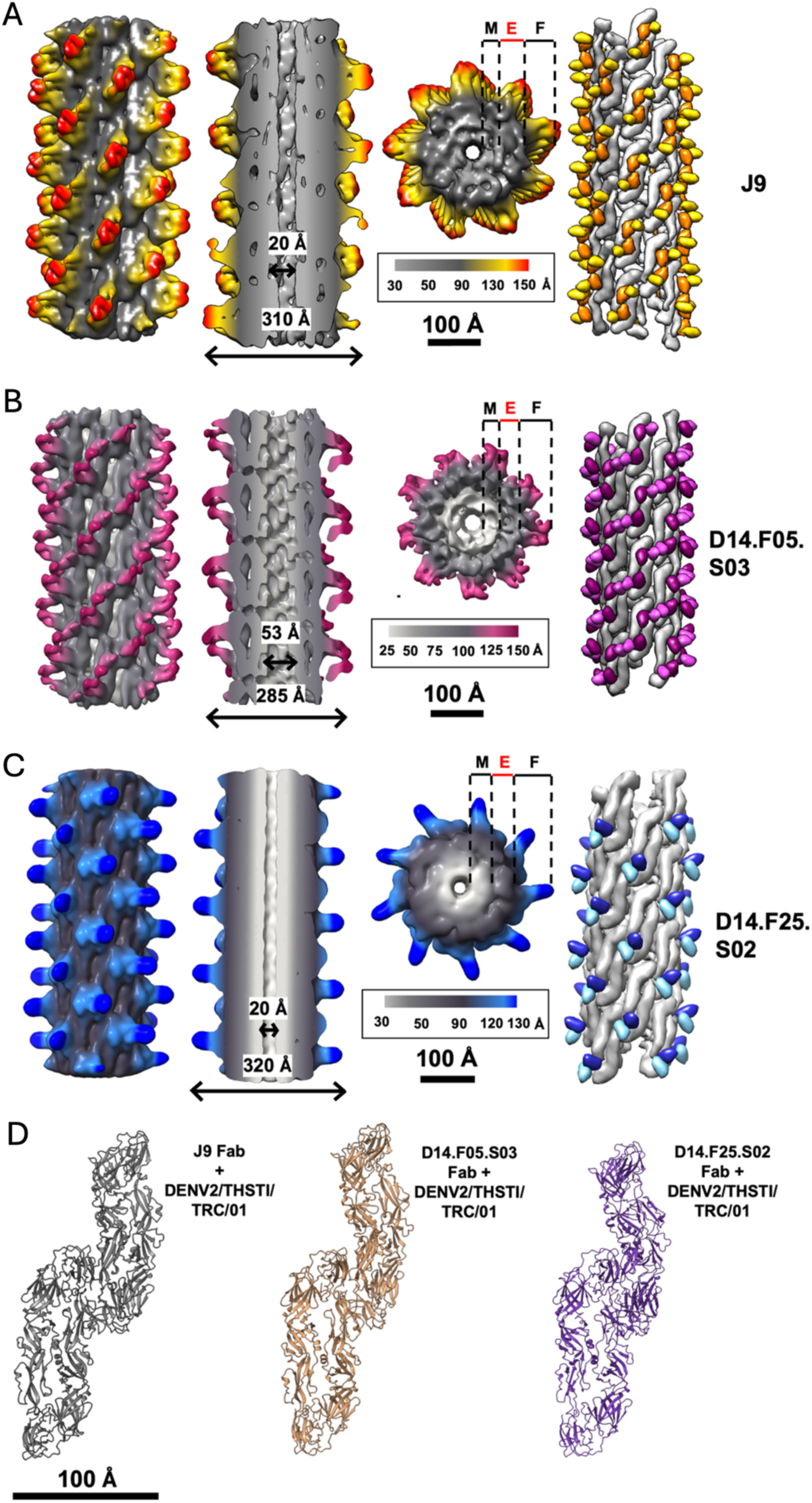
Helical reconstruction of DENV2/THSTI/TRC/01 complexed with J9, D14.F05.S03 and D14.F25.S02 Fab fragments. Side view, central cut-out, top view and surface rendered helical density map of DENV2/THSTI/TRC/01 + J9 Fab (**A**), DENV2/THSTI/TRC/01 + D14.F05.S03 Fab (**B**), and DENV2/THSTI/TRC/01 + D14.F25.S02 Fab (**C**) complexes. Each helical density map is coloured based on its radius from the central helical axis with only the Fab regions shown in contrasting colours. In the surface rendered view (right panels of A-C), only the Fab variable region is shown. **D**. Comparison of adjacent E-dimer positions in all three Fab-bound THSTI/TRC/01 helical structures showing similar E-dimer arrangements. This similarity in adjacent E-dimer arrangement is reflected in the similar helical parameters obtained for the three helical structures of DENV2/THSTI/TRC/01 complexed with J9 (Rise=15.19 Å, Twist=80.62°), D14.F05.S03 (Rise=15.01 Å, Twist=79.94°) and D14.F25.S02 (Rise=15.13 Å, Twist=80.09°).

Despite the differences in tube diameters observed in all three helical THSTI/TRC/01 structures, the E-dimers adopt a similar steep step-like arrangement of E-dimer around the membrane tube (Fig.5). This arrangement differs drastically from the E-dimer arrangement observed in the helical US/BID-V594/2006 structures (Fig.4), demonstrating that despite binding of the same bnAbs, the E-protein arrangement on tubular particles is dictated by the virus strain properties. In both the J9- and D14.F05.S03-bound helical structures from THSTI/TRC/01, two Fabs are bound per E-dimer with the external tips of the Fab densities joined to each other. Rigid body fitting of the Fab atomic models into the map density indicates that the constant regions of the Fabs would clash with each other in this conformation (Fig.5A,B, Figs. S11D,E, S12E-G). However, no such clashes were observed in the respective icosahedral structural models (Fig.1D,F, Figs.S11E and S12F,G). In spherical DENV virions, the E-protein conformation has a slight curve to match the curvature of the virus. Modelling the Fab structures to a flat conformation of the E-proteins, generally observed in purified E-protein structures(*20*, *27*), shows clashing of Fab constant domains similar to that observed in the helical structures of the DENV-THSTI/TRC/01+ Fab complexes (Fig.S11D,E and S12E-G). These analyses suggest that the E-dimer adopts a flatter conformation in the tubular virions when compared to its spherical counterpart. In case of D14.F25.S02, only one Fab density is observed per E-dimer in the DENV-THSTI/TRC/01 tubular reconstruction. Fitting the atomic models of the E-dimers and D14.F25.S02 Fab variable region in the cryo-EM map suggests that the close juxtaposition of the tips of adjacent E-dimers in the helical structure restricts binding of a second Fab due to probable clashes (Fig.5C, Fig.S11F-K).

### D14.F25.S02-binding induced distortion of spherical DENV

Three-dimensional (3D) classification of spherical virions from the DENV2-US/BID-V594/2006 + D14.F25.S02 Fab complex dataset resulted in two distinct classes of bound virions. The larger class of particles contained Fab bound to only the icosahedral 5f and 3f symmetry positions with negligible density observed at the icosahedral 2f position (hereafter referred to as 5f-3f map) (Fig.1E,H). The Fab occupancy is very low in the cryo-EM maps of D14.F25.S02 complex, with 5f and 3f Fab occupancy estimated to be approximately 28 % and 18 % respectively (calculated as percentage of the average map value of Fab/E-protein(*28*)).

The smaller 3D class contained particles where the D14.F25.S02 Fab was bound only at the icosahedral 3f and 2f positions (hereafter referred to as 3f-2f map) (Fig.6A). Refinement of the 3f-2f class, with icosahedral symmetry imposed, resulted in a cryo-EM map of 6.48 Å global resolution (Fig6A, Fig.S14A). Sub-particle, asymmetric 3D refinement of the E-raft resulted in a map of 3.53 Å resolution, where only the 3f-bound Fab had defined density to allow real-space refinement (Fig.S14B,C). Analysis of the icosahedral map shows that the Fab occupancy in the 3f-2f map was 33% and 23% for the 3f and 2f positions respectively. However, only ∼20% of particles formed the 3f-2f class compared to the 5f-3f class, indicating that the Fab binds to the icosahedral 5f position robustly. The Fab structure, footprint and angle of approach are equivalent in both 3f-2f and 5f-3f maps (Fig.1E, 2B,E, Fig.6A-C, Fig.S14C,D). Though there is no evidence of potential clashes, comparing the Fab structural models bound to symmetry-related positions on the virion suggest that binding of D14.F25.S02 to 5f position affects Fab binding to the 2f positions, as is seen in the negligible Fab occupancy at 2f positions in the 5f-3f map. Analysis of 2D classes shows that saturation level binding of D14.F25.S02 bnAb causes distortion of the spherical virions (Fig.S14E). Spherical virions are only observed in 2D classes that contain feeble Fab projections suggesting lower binding ratio of Fab molecules to virions (Fig.S14E).

**Figure 6.**
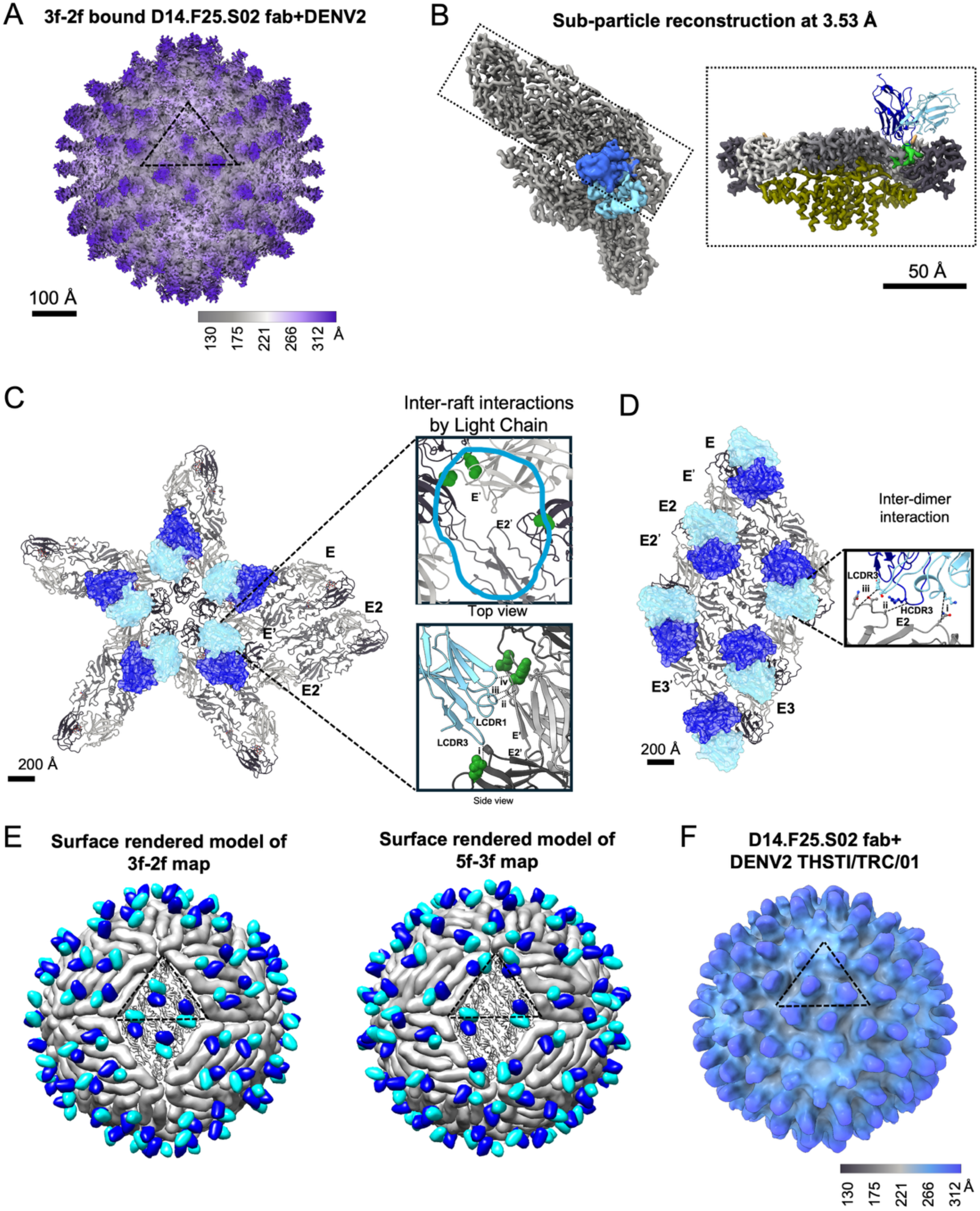
Cryo-EM reconstructions of D14.F25.S02 Fab bound to DENV strains. **A**. Surface rendered 3f-2f cryo-EM density map of D14.F25.S02 Fab + DENV2-US/BID-V594/2006 complex. **B**. Density map cutout from sub-particle reconstruction of 3f-2f map showing E-protein density and bound Fab. Left panel shows the top view of the asymmetric unit with both Fab and E-protein rendered as map density. Right panel shows side view of the E-dimer with Fab complex with trans-membrane regions colored in olive green. **C**. Top view of icosahedral 5f position from D14.F25.S02 Fab + DENV2-US/BID-V594/2006 complex showing inter-E-raft interactions made by D14.F25.S02 light chain. E-proteins are shown as ribbons and Fabs are shown as surface representations. Zoomed-in panels show top view and side view of inter-E-raft interaction regions. Amino acid residues important for neutralization have been depicted as green-colored balls. In the zoomed-in top view, cyan outline indicates the Fab footprint. In the side-view, contacts between the Fab light chain and E-protein residues have been labelled as dotted lines, (i) Y97->P384 (ii) S26->E174 (iii) Y32->G179 (iv) G70->T176 respectively. Residue contacts are calculated as residues with <8Å distance between Ca-Ca backbone of Fab and E-protein respectively. **D**. Top view of inter-E-dimer interactions within an E-raft from the D14.F25.S02 Fab + DENV2-US/BID-V594/2006 complex structure showing inter-E-dimer interactions made by Fab light chain, contacts have been labelled as dotted lines, (i) K33->Q52 (ii) R105->A 224 (iii) Y97->Q227. **E**. Surface rendered model of the 5f-3f and 3f-2f structures of D14.F25.S02 Fab complexed to DENV virion. E-protein is shown in grey and the Fab variable regions are shown in dark blue for heavy chain and light blue for the light chain. **F**. Surface rendered cryo-EM density map of D14.F25.S02 Fab + DENV2-THSTI/TRC/01 complex. Panels A and F are colored radially from the center and icosahedral asymmetric unit labelled as a dotted triangle.

Analysing the binding epitope of D14.F25.S02 shows that the bnAb makes contacts at both intra-E-dimer and inter-E-dimer regions with the Fab light chain primarily involved in inter-E-dimer contacts between adjacent E-rafts (Fig.6C,D and Fig.S8). At the icosahedral 5f position, the Fab light chain has a different local environment when compared to the 2f and 3f positions and makes numerous contacts to the DI and DIII domains of E-proteins in adjacent E-dimer rafts (Fig. 6C, Table S5). E-protein residues, Tyr 178 and Met 297, which have been implicated in neutralization activity of D14.F25.S02(*14*), are located within bnAb footprint at the 5f position, indicating that these inter-raft contacts are essential for the bnAb activity (Fig.6C). In contrast, D14.F25.S02’s interactions with adjacent E-dimers within an E-raft are minimal (Fig.6D, Table S5) and not necessary for its binding or neutralization activity(*14*). These observations further suggest that D14.F25.S02 binds robustly to the 5f position compared to the 3f and 2f positions on the virion (Fig. 6E).

Cryo-EM analyses of DENV2-US/BID-V594/2006 complexed to D14.F25.S02 IgG was also attempted to get a broader sense of bnAb epitope binding (Fig.S15). Massive aggregation of virions was observed in the dataset with few isolated virions (Fig.S15B). Single-particle cryo-EM reconstruction was calculated from a small subset of non-aggregated and loosely aggregated virions to a global resolution of 7.7 Å (Fig.S15). The IgG binding positions on the virion are identical to that observed with the Fab (Figure 1D, 2E, 6B and Fig.S15), and with low binding occupancy (26%) at all symmetry-related epitope sites. 3D classification did not yield any differentially bound IgG subsets as observed with the Fab-bound dataset, which could be due to the lower number of isolated particles in the IgG dataset. The IgG map density could be resolved only till the Fab region, and no density was observed for the Fc portion of the IgG, even at very low thresholds (Fig.S15F). This observation along with massive aggregation seen in the dataset suggests that the two arms of the IgG cross-link adjacent virions. The overall low occupancy of Fab arms in the averaged IgG+DENV complex reconstruction further reinforces that the D14.F25.S02 binding is relatively weak across different epitope positions on the virus.

Despite several attempts, it was not possible to achieve a high resolution cryo-EM structure of D14.F25.S02 Fab complexed to the DENV2-THSTI/TRC/01 strain. The best possible 3D reconstructed map of D14.F25.S02 Fab+DENV2-THSTI/TRC/01 complex achieved a global resolution of ∼19 Å showing Fabs bound equivalently at all symmetry related positions in the cryo-EM map (Fig.6F). However, the Fab and E-protein features lack definition, suggesting higher variability in the constituent virions (Fig.6F, Fig.S16). Analysis of 2D classes from the DENV2-THSTI/TRC/01+ D14.F25.S02 Fab dataset showed clear distortion of Fab-bound virion particles, indicating that the virus morphology was dramatically affected by D14.F25.S02 Fab binding (Fig.S16B).

The highest resolution cryo-EM structure of J9+DENV was achieved in complex with the DENV2-US/BID-V594/2006 strain (Fig.1D,G). In case of D14.F05.S03, high resolution structure determination was achieved in complex with the DENV2-THSTI/TRC/01 strain (Fig.1F,I). Additionally, a 7.2 Å cryo-EM structure of J9 Fab complexed with DENV2-THSTI/TRC/01 was also determined, which showed clear density for the J9 Fab and identical binding features as seen with the DENV2-US/BID-V594/2006 strain (Fig.S17). These results demonstrate that the DENV2-THSTI/TRC/01 strain consists of homogenous mature particles suitable for single-particle cryo-EM analyses. These results also highlight that it is the binding of D14.F25.S02 Fab to DENV2 and not DENV2-strain dependent properties which impeded higher resolution cryo-EM structure determination of D14.F25.S02 Fab-bound DENV2-THSTI/TRC/01 sample.

Furthermore, in comparison to the D14.F25.S02 bound spherical DENV structures, the occupancy of bound Fab was much higher in the case of D14.F05.S03 and J9. D14.F05.S03 showed high occupancy of Fabs at its 3f and 2f positions (82%) with relatively lower occupancy (50%) at the 5f positions (Fig.1F). In case of J9, the fab bound equivalently to the 5f and 3f positions (∼60%) with lower occupancy at the 2f position (∼35%) (Fig.1D). Analyses of fab interactions using the real-space refined atomic models show that both D14.F05.S03 and J9 make minimal yet similar inter-E-dimer and inter-E-raft contacts (Table S4, S6), with none of the involved residues identified as essential for neutralization activity of these two bnAbs.

### Correlation between binding affinity and neutralization potency of HEDR bnAbs

To evaluate the binding of HEDR bnAbs to DENV and the role of E-dimer as the binding unit, biolayer interferometry (BLI) was used to determine binding kinetics of the three HEDR bnAbs with purified E-dimer ectodomains from all four DENV serotypes (Fig.7). All three HEDR bnAbs show binding affinity in the low nanomolar (nM) range to the E-dimer, exhibiting strong binding and further emphasizing the E-dimer’s role as the primary binding unit. In case of D14.F25.S02, the binding to DENV2 E-dimer is much weaker compared to other DENV serotypes. D14.F25.S02 also shows weaker binding to DENV2 when compared with J9 and D14.F05.S03. These results corroborate with the low occupancy of D14.F25.S02 observed on the DENV2 virion in our cryo-EM analyses. Furthermore, the binding analyses show that the binding affinity of D14.F25.S02 to DENV serotypes 1-4 (Fig.7) follows a similar trend as its reported neutralization potencies (*14*), indicating a direct correlation between the bnAb’s E-dimer binding affinity with virus neutralization potency.

**Figure 7.**
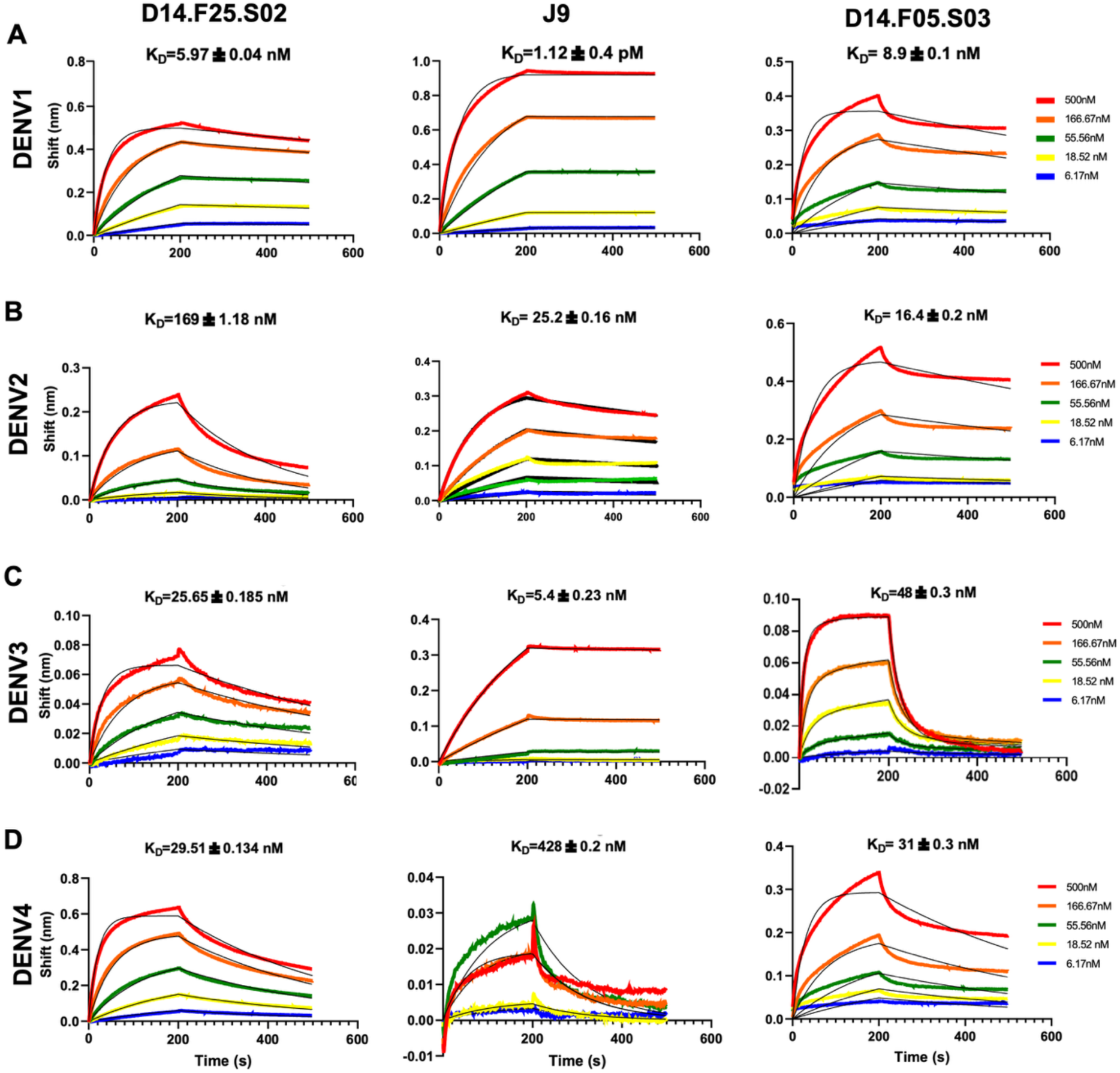
Binding kinetics of HEDR bnAbs to DENV E-dimer. IgG1 binding kinetics of D14.F25.S02 (left column), J9 (middle column) and D14.F05.S03 (right column) to stabilized E-protein dimers from DENV 1-4 serotypes (top to bottom rows). Biolayer interferometry (BLI) sensograms at different concentrations of the analyte with theoretical fitted curves (black) are shown. Analyte concentrations are shown in different colours as per the panel legend. Calculated dissociation constants (K_D_) for each sample complex are also given.

## Discussion

HEDR bnAbs bind mature as well as tubular DENV as shown in this work, along with incompletely mature virions as reported previously(*14*, *15*). Expanded/bumpy DENV particles have been described to be present only at 37°C or above (*24*, *25*) and all three bnAbs described here potently neutralize DENV at 37°C (*14*, *15*). The different DENV morphologies, ranging from spherical (mature, partially mature, immature), to expanded and tubular virions, present varying quaternary surface epitope arrangements and topology. Our current work along with previous reports(*29*) shows that bnAb binding is differential to different DENV morphologies, suggesting that these variation in DENV morphologies might serve as an adaptive mechanism for escaping antibody-mediated neutralization and immune evasion by the virus. The common surface feature across these virion forms is the presence of the E-dimer configuration, which further emphasizes the importance of the E-dimer quaternary unit as a critical antigenic target.

On comparing the five tubular DENV structures reported here, the helical arrangements of E-dimers were similar within each DENV strain type, irrespective of the Fab bound. The binding of an antibody, that locks the E-dimers together, seems to be a prerequisite to form a globally regular arrangement of E-dimers on tubular DENV, but the specific arrangement achieved by the E-dimers is influenced by the virus strain type. In case of DENV2-US/BID-V594/2006 strain, the helical arrangement resembles the E-raft sequentially wrapping around the tubular virion’s length, suggesting tighter inter-E-dimer and inter-E-raft interactions. However, in the THSTI/TRC/01 strain, the relative strength of inter-E-dimer and E-raft interactions appears to be weaker as the helical arrangement is drastically different from the icosahedral E-raft arrangement.

The HEDR bnAb complex structures described here, show that these bnAbs occupy similar epitopes that bridge the E-dimer and mask the fusion loop, strongly suggesting that the bnAbs lock the E-dimers in the pre-fusion state. The global bnAb footprints overlap with prM binding region on E-protein, which is similar to that observed with EDE bnAbs (*17*, *20*), highlighting the conserved nature of this epitope. Fusion loop exposure on the E-protein is necessary for attachment to endosomal membranes and to initiate viral membrane fusion in dengue virus cellular entry(*5*, *6*). Similarly, dissociation of E-dimers is necessary for conformational changes in the E-proteins from pre-fusion to post-fusion structure, which has again been shown to be essential for successful virus infection in cells(*30*, *31*). Thus, the bnAbs likely inhibit virus infection in cells by blocking exposure of the fusion loop and impeding E-dimer dissociation, both steps that are essential for viral entry into host cells. Indeed, previous report on the J9 bnAb has shown that the bnAb obstructs early entry steps in virus infection(*15*), which further corroborates our results.

The heavy chain of the HEDR bnAbs make majority of the contacts within an E-dimer, including the b-strand and fusion loop region. The involvement of the heavy chain is also predominant in interactions with the conserved glycans and makes robust contacts than the respective Fab’s light chains. The light chain of the bnAbs is involved in inter-E-dimer contacts both within and between E-rafts, likely stabilizing the bnAb binding on the virions. As seen in case of D14.F25.S02, the inter-E-raft light chain interactions are specific and necessary for neutralization activity of the bnAb. However, the bnAbs also bind the tubular forms of DENV with good occupancy despite the disruption in bnAb light-chain contacts as evident in the helical DENV2-THSTI/TRC/01 complex structures. Thus, the bnAb heavy chain drives the overall epitope recognition and potency of these HEDR antibodies.

From our results, it is evident that binding of HEDR category 1 bnAb, D14.F25.S02, distorts spherical DENV, whereas the HDER category 2 bnAbs, J9 and D14.F05.S03 show no such effects, despite the overall similar binding features shared between the three bnAbs. Though the epitope features of D14.F25.S02 on the E-dimer are equivalent across the multiple virion structural forms determined in this study, and the recently reported structure of D14.F25.S02 Fab bound to purified DENV3 E-dimer ectodomain (*32*), our results demonstrate that the antibody binding is affected by the virion topology. In case of the DENV2-THST/TRC/01 strain, the effect of D14.F25.S02 binding at saturating concentrations has a dramatic effect on virion structure and symmetry, impeding higher resolution structure determination of the virion+Fab complex. However, the Fab binding did not destroy virion organization but rather appears to have introduced significant local movements in the E-proteins, as the virions’ icosahedral symmetry was still discernible from the low-resolution 3D density map. Remarkably, a high resolution cryo-EM structure was achievable for D14.F25.S02 to DENV2-US/BID-V594/2006 strain, indicating that this strain was relatively resistant to distortion due to bnAb binding which could be a further reflection of stronger interactions between the E-dimer and E-rafts in the US/BID-V594/2006 strain when compared to THSTI/TRC/01 strain. Nevertheless, analyses of 2D classes and cryo-EM density maps show that binding of D14.F25.S02 to the US/BID-V594/2006 virions does affect the topology of the E-protein surface, leading to many virion particles being discarded during the 3D reconstruction process.

The major class of particles obtained from the D14.F25.S02+DENV2-US/BID-V594/2006 dataset had the highest Fab occupancy at the icosahedral 5f position. The local E-dimer arrangement at the 5f position likely facilitates D14.F25.S02 binding as the larger gap between E-dimers surrounding the 5f vertices would allow easier insertion of the bnAb HCDR3 loop between adjacent E-dimers. The bnAb also makes inter-raft contacts that are unique to the 5f position which have been reported to be crucial for its neutralization activity. These observations indicate that D14.F25.S02 binds preferentially to the 5f position on the icosahedral virus. However, the binding at 5f vertices effectively inhibits bnAb binding to the icosahedral 2f position, suggesting that the bnAb binding at 5f causes local structural variations which limits bnAb binding at the other sites. Notably, when D14.F25.S02 binds the 2f and 3f positions on the virus, there is negligible binding observed at the 5f position, suggesting that once D14.F25.S02 binds at 2f position, bnAb binding at 5f position is hampered. These observations imply that binding of D14.F25.S02 at the 5f position causes perturbations in the adjacent E-raft which impedes bnAb binding to the 2f position, whereas binding of bnAb first to the 2f and 3f positions stabilizes the E-raft such that E-dimer movement near the 5f position is restricted and thus, could inhibit bnAb binding. Similar reports of local distortion to DENV structure have been reported for EDE1 bnAb C10 which also strongly neutralizes DENV and ZIKV(*28*). Binding of C10 to purified DENV E-dimers has been shown to cause hinge-like motions in the E-protein(*17*), along with preferred binding of bnAb to 5f and 3f positions and increased dynamics of E-proteins with increasing Fab concentrations(*28*). Thus, despite differences in binding determinants and specific interactions to the E-proteins, HEDR category-1 bnAbs and EDE1 bnAbs show similarities in disrupting E-protein organization on spherical DENV surface.

Structural elucidation of these pan-DENV bnAb epitopes, as described here, emphasizes the need for vaccine immunogen design that most closely imitate the E-protein quaternary epitopes, including glycosylation, as presented on mature DENV surface to elicit broad neutralization responses. For both, D14.F25.S02 and D14.F05.S03, bivalent engagement as IgGs is necessary for potent neutralization activity against DENV(*14*). Based on the structural data obtained from D14.F25.S02 IgG bound to DENV2 and previous published reports(*13*, *33*, *34*), it seems possible that virion aggregation along with locking of E-dimers maybe essential for the neutralization activity of these bnAbs. Despite the distortion caused by D14.F25.S02 binding to DENV, the bnAb still has one of the highest reported neutralization potencies for a pan-DENV and ZIKV bnAb(*14*). When comparing neutralization of DENV serotypes alone, J9 has one of the highest reported neutralization potencies reported till-date(*15*). J9 can neutralize DENV as IgG as well as Fab fragment. The potent activity of J9 as Fab fragment increases its applicability as a therapeutic antibody while avoiding any ADE effects caused by Fc regions present on IgGs. Similarly, despite the need for bivalent engagement, antibody engineering (*35, 36*) of D14.F25.S02 could be leveraged for developing the bnAb as a pan-DENV+ZIKV therapeutic antibody. Thus, beyond expanding fundamental insights into pan-serotype epitope recognition by bnAbs targeting DENV, our findings also highlight the potential of engineered bnAb variants as therapeutic options for severe dengue infections.

## Supporting information

all supplemental material combined

## Acknowledgments

We thank the Advanced Center for Cryo-Electron Microscopy (ACCEM) facility at Indian Institute of Science, Bangalore (IISc) and the DST-SATHI Cryo-EM facility at IIT-Delhi for data collection time and support. We acknowledge the Department of Biotechnology, Department of Science and Technology (DST), and Ministry of Human Resource Development, India for funding the cryo-EM facility at IISc-Bangalore. We acknowledge DBT-BUILDER Program (BT/INF/22/SP22844/2017) and DST-FIST (SR/FST/LSII-039/2015) for the National Cryo-EM facility at IISc, Bangalore. We thank the Bioimaging facility at IISc which houses the 120kV room temperature TEM for time and support. The authors also thank the x-ray facility at Molecular Biophysics Unit, IISc, funded by DST-SERB grant IR/SO/LU/0003/2010-PHASE-II for loaning dry-shipper for sample shipments to cryo-EM facilities. We thank Mr. Susheelendra Vaidya, FAS-Octet Technical Expert Asia Pacific and Mr. Manish Nag from Prof. Raghavan Vardarajan’s lab at IISc for advice and assistance with the BLI experiments. We also thank Dr. Kelly Lee, University of Washington for his support during project initiation and scientific discussions.

## Author contributions

Conceptualization: L.G and V.M.P

Methodology: A.C, A.R, S.S, S.C, J.L

Investigation: A.C, A.R, S.S, V.M.P

Funding acquisition: V.M.P

Supervision: V.M.P

Writing, V.M.P, A.C, A.R and S.S

## Data availability

Cryo-electron microscopy density maps and corresponding atomic models presented in this study has been submitted to the Electron Microscopy Data Bank (EMDB) and RCSB Protein Data Bank (PDB). Accession codes for the same are as follows:

| S.no. | Sample description | EMDB<br>ID | PDB<br>ID |
| --- | --- | --- | --- |
| 1. | J9 Fab + DENV2-US/BID-V594/2006 icosahedral reconstruction | EMD-63759 | 9MAV |
| 2. | Sub-particle reconstruction of J9 Fab + DENV2-US/BID-V594/2006 icosahedral reconstruction | EMD-80279 | 25OV |
| 3. | J9 Fab + DENV2-US/BID-V594/2006 helical reconstruction | EMD-63762 | 9MAZ |
| 4. | J9 Fab + DENV2-THSTI/TRC/01 icosahedral reconstruction | EMD-63763 | 9MB0 |
| 5. | J9 Fab + DENV2-THSTI/TRC/01 helical reconstruction | EMD-63764 | 9MB1 |
| 6. | D14.F25.S02 Fab + DENV2-US/BID-V594/2006 icosahedral reconstruction (5f-3f map) | EMD-63643 | 9M5H |
| 7. | Sub-particle reconstruction from 5f-3f icosahedral map | EMD-80390 | 25US |
| 8. | D14.F25.S02 Fab + DENV2-US/BID-V594/2006 icosahedral reconstruction (3f-2f map) | EMD-63813 | 9U38 |
| 9. | Sub-particle reconstruction from 3f-2f icosahedral map | EMD-80426 | 25WK |
| 10. | D14.F25.S02 IgG + DENV2-US/BID-V594/2006<br>icosahedral reconstruction | EMD-63812 | 9U37 |
| 11. | D14.F25.S02 Fab + DENV2-US/BID-V594/2006 helical<br>reconstruction | EMD-63809 | 9U34 |
| 12. | D14.F25.S02 Fab + DENV2-THSTI/TRC/01 helical<br>reconstruction | EMD-63810 | 9U35 |
| 13. | D14.F25.S02 Fab + DENV2-THSTI/TRC/01<br>icosahedral reconstruction | EMD-63811 | 9U36 |
| 14. | D14.F05.S03 Fab + DENV2-THSTI/TRC/01<br>icosahedral reconstruction | EMD-80506 | 26BC |
| 15. | Sub-particle reconstruction of D14.F05.S03 Fab +<br>DENV2-THSTI/TRC/01 | EMD-80540 | 26CL |
| 16. | D14.F05.S03 Fab + DENV2-THSTI/TRC/01 helical<br>reconstruction | EMD-80544 | 26CR |

## Supplementary Materials

### Materials and Methods

Figs. S1 to S17

Tables S1 to S7

References (1–21)

