## Supplementary material for "Structural basis of broadly neutralizing human antibodies targeting diverse dengue virus forms": all supplemental material combined

### Materials and Methods

Virus sample preparation:

Dengue virus serotype 2 strains THSTI/TRC/01 (GenBank: UNZ79673.1) was obtained as a kind gift from Dr. Guruprasad Medigishi, THSTI, Faridabad and US/BID-V594/2006 (GenBank: ACA48992.1) was obtained through BEI Resources, NIAID, NIH: Dengue Virus Type2, DENV-2/US/BID-V594/2006, NR-43280. Both virus strains were propagated with minimal passages in mosquito cells (P-6) to 1L culture volumes and purified in a similar manner as reported previously(*1*). Mosquito cells (C6/36) were cultured in Dulbecco's Modified Eagle Medium (DMEM) supplemented with 10% fetal bovine serum (FBS) and 1% nonessential amino acids (NEAA). The cells were maintained at 28°C with 5% CO2 in a 10-stack cell factory system (Nunc). Upon reaching 70-75% confluency, cells were infected with DENV-2 using a multiplicity of infection (MOI) of 0.1. The infection was allowed to proceed at 30°C with 5% CO2. Post 2 hours of infection, the media was changed to complete media supplemented with 5% FBS. At 20 hours post-infection, media was again changed to complete media supplemented with 2% FBS. Supernatant medium containing the virus was collected on 4th and 6th day post-infection. Virus-containing media was centrifuged at 9500xg for 30 minutes and filtered through a 0.22-micron filter to remove all cell debris. The filtrate was precipitated with 8% polyethylene glycol (PEG) overnight at 4°C, followed by centrifugation at 9500xg for one hour to collect the PEG pellet. The pellet was then resuspended in NTE buffer (120 mM NaCl, 20 mM Tris, 1 mM EDTA, pH 8.0). The resuspended PEG pellet was sedimented through a 24% sucrose cushion. The dissolved sucrose pellet applied to a potassium tartrate gradient (35%-10%) and centrifugated at 32000 rpm for 2 hours at 4°C in a SW41-Ti swinging bucket rotor (Beckman Coulter). Bands corresponding to the virus was extracted from the 20% fraction of the discontinuous gradient, buffer exchanged and concentrated in NTE buffer using a 100 kDa centrifugal filter. The purity and concentration of the virus sample was estimated by visualizing under negative staining transmission electron microscopy (TEM) analysis.

Fab preparation:

Purified IgG of D14.F25.S02, J9, and D14.F05.S03 bnAbs were prepared as previously reported(*2*, *3*). BnAbs were digested with papain individually to generate Fab fragments (~50kDa molecular weight) using a commercially available Fab Fragmentation kit (G Biosciences) (Cat. # 786-273). Fabs were then separated in flowthrough by passing over a Protein-A column. Concentrations of Fabs obtained for F25.S02, J9, and F05.S03 were 0.9 mg/ml, 0.6 mg/ml and 1.01 mg/ml, respectively. Concentration of F25.S02 IgG sample was 1mg/ml.

E-dimer expression and purification:

Stabilized E-dimer ectodomain plasmid constructs were a gift from Dr. Brian Kuhlman, obtained via Addgene (plasmid # 219718, 219719, 219720, 219721). The plasmid constructs were expressed in Expi293 suspension cell lines and purified according to previous reports(*4*). The purified protein constructs were confirmed for presence of dimers via size-exclusion chromatography and negative stain transmission electron microscopy (nsTEM).

Cryo-EM sample preparation:

Purified DENV samples at an approximate concentration of 0.25 mg/ml (E-glycoprotein concentration) were incubated with the respective Fabs and F25.S02 IgG at room temperature for 45 minutes. In case of D14.F25.S02, molar ratio of Fab to virus was 4:1 and molar ratio of IgG to virus was 1:1. For J9 and D14.F05.S03, molar ratio of Fab to virus was maintained at 2:1 and 2.5:1 respectively. For estimating the concentration of dengue virus needed for the experiments, the concentration of E-glycoprotein estimated from SDS-PAGE was used. Ultra-thin lacey carbon grids (300 or 400 mesh) were glow discharged using 25 mA current for 30s with a PELCO EasiGlow machine. Aliquots of 3ul virus-Ab mixture were applied on glow discharged grids and plunge-frozen with FEI Vitrobot mark IV for DENV-2-THST1/TRC/01 + Fab complexes (Blot time – 8s, Temperature - 4° C, Humidity – 100%) and Leica EM GP2 for DENV2-US/BID-V594/2006 + Ab complexes (Blot time – 3.5s, Temperature - 4° C, Humidity – 100%).

Cryo-EM data collection:

All DENV-2-THST1/TRC/01 and Fab complex samples were imaged in Talos Arctica (FEI, ThermoFisher Inc.), operating at 200 kV, with a K2 summit electron detector (Gatan Inc.). Micrographs were collected with a pixel size of 1.16 Å/pixel, at a total dose of 38 - 40 $e^-/Å^2$ fractionated into 20 frames per image, and with defocus range of -1.5 to -2.5 µm.

DENV2 US/BID-V594/2006 samples complexed with J9 Fab, F25.S02 Fab and IgG were imaged using Titan Krios G3I, operating at 300 kV with a K3 detector and an energy filter (Bioquantum imaging) operating at a slit width 20eV. Micrographs for each dataset were collected with a total dose of 40 – 42 $e^-/Å^2$ fractionated into 38 frames per image at a defocus range of -1 µm to -2.5µm and with a pixel size of 1.1 Å/pixel.

Cryo-EM data processing:

A combination of RELION-4.0 (*5*) and cryoSPARC version 4.7.0 software package (*6*) was used for data processing following standard single particle analysis (SPA) steps. Frame alignment was carried out using Relion's own implementation of motion correction or Cryosparc's Patch motion correction. CTF estimation was performed using CTFFIND4 (*7*).

J9-virus complex datasets: A total of 4756 micrographs were collected for the J9 + DENV2 US/BID-V594/2006 complex at a magnification of 81000x with corresponding pixel size of 1.1Å/Pixel. Around 15000 particles were manually picked in RELION-4.0. Further, the particle coordinates were imported to cryoSPARC-4.7.0. After 2D and 3D classification, 2900 best particles were used for iterative non-uniform refinement and CTF refinement, resulting in an electron density map of 3.85 Å global resolution.

After full virus icosahedral reconstruction, sub-particle reconstruction was performed in cryoSPARC-4.7.0 using its local reconstruction pipeline. For local reconstruction, at first, symmetry expansion was performed with a particle set from the initial 3D-refinement after the first 3D-classification, consisting of 3577 full icosahedral particles. Total 2,14,620 sub-particles were recentered to the virus asymmetric unit axis and re-extracted with applied shifts in coordinates. A smooth, soft, static mask was generated from an asymmetric unit cutout of the virus. Using this mask, an initial focused 3D-classification was performed from which 3D-classes with clear density of fab was selected for further processing. With the selected classes and static mask, initial local refinement along with CTF refinement was performed which yield a 3D reconstruction of resolution 3.3 Å. Further, local refinement was done with dynamic mask which was centered at C1 axis of particle covering a region larger than an E-raft. Using the same mask input, CTF refinements and reference-based motion correction yielded reconstruction of 3Å. A final round of local refinement with dynamic mask which included only the E, M proteins and J9 fab variable region was used which yielded a 3D reconstruction of 2.83Å.

For the J9 + DENV-2-THST1/TRC/01 complex, 1100 images were collected at a magnification of 36000x with corresponding pixel size of 1.16Å/Pixel. Of the 10108 manually picked particles, only 668 particles showed clear Fab densities after 2D classification in RELION-4.0. These particles were then used non-uniform refinement in cryoSPARC-4.7.0. The 3D refinement resulted in an electron density map of 7.21 Å global resolution.

D14.F25.S02-virus complex datasets: For the D14.F25.S02 Fab + DENV2 US/BID-V594/2006 complex, 18,787 micrographs were collected at a magnification of 81000x with corresponding pixel size of 1.1Å/Pixel. Dataset for D14.F25.S02 IgG + DENV2 US/BID-V594/2006 was also collected at the same magnification and microscope settings as the Fab datasets with a total of 4172 images. Virion particles were manually picked from both datasets, with 46,825 particles picked from the Fab dataset and 8176 particles from the IgG dataset. Subsequent rounds of 2D classification and 3D classification were performed on the individual particle datasets. For the Fab complex, a 3D class with 2305 particles was selected and used for non-uniform refinement(*8*) and CTF refinement(*9*) which resulted in a 4.09 Å density map showing Fab density near 5f and 3f symmetry vertices. A smaller 3D class containing 1559 particles was selected for non-uniform refinement which resulted in a 6.48 Å density map of the Fab bound majorly near the 3f-2f symmetry vertices. For the IgG dataset, 2D classification, 3D refinement and post-processing was carried out in RELION-4.0. The final 3D reconstruction of 7.65 Å global resolution was calculated from 1045 particles selected from 3D classification.

To improve resolution and map quality to allow model building, sub-particle reconstruction was performed in cryoSPARC-4.7.0 using its local reconstruction pipeline. Symmetry expansion was performed with the particle set from the final 3D-refinement of the icosahedral virus bound with fab, consisting of 2300 (5f-3f map) and 1559 (3f-2f map) virion particles. A smooth, soft, static mask was generated from a raft cutout of the virus. Total 138,300 and 93,540 sub-particles respectively from 5f-3f and 3f-2f map, were recentered to the axis of the mask and re-extracted with applied shifts in coordinates. Initial focused 3D-classification was performed using the generated focus mask from which 3D-classes with clear density of fab were selected for further processing. With the selected classes and static mask, initial local refinement along with CTF refinement was performed. Further, local refinement was done with dynamic mask which was centered at C1 axis of particle covering a region larger than an E-raft. Using the same mask input, CTF refinements and reference-based motion correction yielded reconstruction of 3.19Å and 3.78 Å respectively for 5f-3f and 3f-2f map. A final round of local refinement with dynamic mask which included only the E and M proteins and D14.F25.S02 fab variable region was used which yielded a 3D reconstruction of 3.08Å and 3.53 Å respectively for 5f-3f and 3f-2f map.

For the D14.F25.S02 Fab + DENV-2-THST1/TRC/01 complex, 4000 micrographs were collected at a magnification of 36000x with corresponding pixel size of 1.16Å/Pixel. From these images, 26782 particles were manually picked and subjected to unsupervised 2D classification in RELION-4.0. The best classes which showed apparent Fab densities were selected for reference-based 3D classification in RELION-4.0. Different starting models, including a sphere, mature

DENV structure and the determined DENV2-US/BID/V594/2006+D14.F25.S02 Fab structure were used for 3D classification. All the classification routines resulted in similar results. Finally, a total of 4715 particles from the best classes of 3D classification from RELION4 were utilized for 3D non-uniform refinement in cryoSPARC-4.7.0 to generate a 3D density map at 19 Å global resolution.

D14.F05.S03-virus complex datasets: A total of 3176 micrographs were collected at a magnification of 36000x with corresponding pixel size of 1.16Å/pixel for D14.F05.S03 + DENV-2-THST1/TRC/01 complex. From these images, 32650 particles were manually picked using RELION-4.0 software. The manually picked particle coordinates were imported to cryoSPARC-4.7.0 for further processing. A final set of 4299 particles were selected for 3D refinement following non-reference 2D classification and reference-based 3D classification. Subsequent iterative non-uniform refinement and CTF refinement resulted in a 3D density map with global resolution of 4.51 Å. Sub-particle reconstruction for the icosahedral virion complex map was carried out with similar protocol as described above for the D14.F25.S02 fab complex. Following the local reconstruction process in cryoSPARC-4.7.0 resulted in a 3.1 Å map of the E-raft with Fab variable region.

Real space refinement and model building:

The J9 fab complex map obtained from sub-particle reconstruction was sharpened with DeepEMhancer (*10*). Similarly, the highest resolution sub-particle reconstruction maps of D14.F25.S02 and D14.F05.S03 fab complexes was sharpened using EMReady2 (*11*). With the sharpened maps, de novo model building was performed using EMProt-1.2 (*12*). The output models were checked and corrected for sequence similarity and completeness. In case of D14.F25.S02 fab, the Fab variable region was predicted using AlphaFold3 web server (*13*). The corrected models for all fab complexes (respective fab variable regions and E-proteins) were then taken for real-space refinement in the Phenix software package (*14*, *15*). Only the E-dimer and 5f fab was used for real space refinement in the 5f-3f sub-particle map. Similarly, only the E dimer and 3f fab position was used for real space refinement in the 3f-2f sub-particle map. For all high resolution maps, geometry restraints and Ramachandran clashes were adjusted as needed using COOT (*16*) and ISOLDE (*17*). Glycan components were added to the E-protein atomic models using other higher-resolution E-protein x-ray crystallography structures (PDB IDs: 4UT6, 4UTA) as references. Glycan residues (N-acetyl glucosamine (NAG), α-D-mannopyranose (MAN), α-L-fucopyranose and β-D-mannopyranose (BMA) were manually added to N67 and N153 residues of the E-protein using COOT according to the map densities and were included in the atomic models for real-space refinement. For full virus Fab-complex maps, models obtained from the subparticle refined maps were used for rigid body fitting.

For delineating interactions between fabs and E-protein, contact distances where generated in UCSF ChimeraX (*18*) with an all atom distance cutoff of <5Å for J9 and D14.F05.S03 complex models and <8 Å Cα-Cα distance cutoff for D14.F25.S02 complex model. Maps and models were analyzed and figures prepared using UCSF Chimera(*19*) and UCSF ChimeraX(*18*).

<u>Helical reconstructions of tubular DENV regions:</u>

For the helical reconstruction of tubular regions from DENV2 US/BID-V594/2006 virions, helical segments were manually picked in segment picking of RELION-4.0. For both D14.F25.S02- and J9-bound virions, the initial helical rise and twist for DENV2 US/BID-V594/2006 tubular particles were estimated manually in UCSF Chimera by arranging the Fab+E-dimer models around a cylinder of same dimensions as the virion segments with minimal clashes. A 2D projection of the manually arranged helical geometries was then used to match with the arrangement observed in 2D class averages of tubular particles with no symmetry applied. An initial rise and twist for D14.F25.S02-bound segments (rise=10 Å and twist-=30°) and for J9-bound segments (rise=13Å and twist=36°) was used for reference-free helical 2D classification(*20*) in RELION-4.0. Best 2D classes containing a total of 2188 segments for D14.F25.S02-bound dataset and 1361 segments for J9-bound dataset were subjected to 3D classification with initial estimates using a 3D-cylinder as starting model. In case of D14.F25.SO2 dataset, 405 segments were imported for final helical 3D reconstruction in cryoSPARC-4.7.0 using non-uniform refinement option with rise and twist of 11.25Å and 29.48° which yielded a density map of global resolution 14.7 Å. For the J9 dataset, a best class with 253 segments was used for helical 3D reconstruction in RELION-4.0 with rise and twist of 12.83Å and 35.83° that resulted in a 3D map with estimated global resolution of 44Å.

For bnAb bound tubular-shaped particles found in DENV-2 THSTI/TRC/01 datasets, the tubular-shaped regions of the virions were manually picked following the helical particle picking protocol of RELION-4.0 and unsupervised 2D classification was performed. Class averages with distinct Fourier amplitude patterns were imported to PyHi Fourier Bessel indexing software(*21*) and helical rise and twist values were calculated for the antibody-virus complexes. From the calculated rise and twist, another round of unsupervised helical 2D classification was performed, and the particles were extracted with the same parameters for reference-based 3D classification. For initial 3D classifications 3D-cylinder was used as startup model. The best 3D classes were selected, and the particle co-ordinates were transferred to cryoSPARC-4.7.0 for final helical reconstruction using non-uniform refinement option.

For the J9 + DENV-2 THSTI/TRC/01 complex, 19792 segments were manually picked, and after classification routines, 5769 segments were used for the final 3D refinement, which resulted in a helical density map of 14.41 Å. Similarly, for the D14.F25.S02 and D14.F05.S03 complexes, 8264 and 8045 segments were respectively picked in manual mode, and final sets of 1355 (for D14.F25.S02) and 1311 segments (for D14.F05.S03) were used to generate 3D maps at global resolutions of 17.37 Å and 17.29 Å respectively.

In all helical reconstructions, a hollow cylindrical mask was used which focused on the E-protein and membrane layer of DENV tubular particles during all the refinement steps. Atomic models of the E-protein+fab complexes from the icosahedral virus reconstructions were used for rigid body fitting and interpretation of all the helical reconstruction maps.

<u>Inner volume calculations for capsule-shaped DENV2-US/BID-V4594/2006:</u>

For D14.F25.S02-bound virions:

Average length of cylindrical portion in capsule-shaped DENV = 79 nm

Inner radius of capsule = 5.5 nm.

Thus, internal volume of cylindrical portion in capsule-shaped DENV = 7.503 x $10^{\wedge-24}$ $m^3$

Assuming, volume of one copy of viral genome(*22*, *23*) = 3.8 x 10-24 $m^3$

Number of genome copies that can be possibly accommodated in capsule-shaped DENV = 1.9

For J9-bound virions:

Average length of cylindrical portion in capsule-shaped DENV = 74.149nm

Inner radius of capsule= 6nm

Thus, internal volume of cylindrical portion of capsule-shaped DENV = 8.38 x 10^ $^{-24}$ m3

Assuming, volume of one copy of viral genome(*22*, *23*) = 3.8 x 10-24 $m^3$

Number of genome copies that can be possibly accommodated in capsule-shaped DENV = 2.2

Bio-layer interferometry:

The binding kinetics of three IgGs with E-dimers of DENV serotype 1-4 were determined via BLI on a Sartorius Octet R8e machine. Anti-Penta-HIS capture biosensors (HIS1K) were presoaked in binding buffer phosphate-buffered saline (PBS pH 7.4) supplemented with 0.02% Tween 20 for 15 min. The hydrated tips were then loaded with purified E-proteins prepared at 5 µg/mL in binding buffer for 60 s. D14.F25.S02, D14.F05.S03 IgGs were diluted to 500 nM in buffer and serially diluted three-fold for a final concentration of 6.17 nM in the wells. After reaching a stable baseline, antibody-immobilized biosensors were moved into wells containing the three-fold dilution series of IgGs to monitor association for 200 sec, then biosensors were moved back into wells containing binding buffer to monitor dissociation for 300 sec. Responses were calculated and double-referenced to the buffer reference signal. For J9 IgG and DENV1, 3 and 4, ProG biosensors were used to capture IgG and DENV E-proteins were serially diluted three-fold in the wells. Data acquisition was performed in two independent experiments for each sample complex. Despite multiple attempts, J9 binding to DENV4 E-dimer was relatively noisy compared to the other complexes. Kinetic data were analyzed by using Octet BLI Discovery 13.1.0.25 software and were by processed by Savitzky-Golay filtering prior to fitting using a 1:1 binding model or 2:1 binding model (only for D14.F05.S03+DENV3 sample).

```
DV2 16681 : MRCIGMSNRDFVEGVSGGSWVDIVLEHGSCVTTMAKNKPTLDFELIKTEAKQPATLRKYC     60
DV2 US-BID: MRCIGISNRDFVEGVSGGSWVDIVLEHGSCVTTMAKNKPTLDFELIKTEAKQPATLRKYC     60
DV2 THSTI : MRCIGISNRDFVEGVSGGSWVDIVLEHGSCVTTMAKNKPTLDFELIKTEAKQPATLRKYC     60
            *****:******************************************************

DV2 16681 : IEAKLTNTTTESRCPTQGEPSLNEEQDKRFVCKHSMVDRGWGNGCGLFGKGGIVTCAMFR     120
DV2 US-BID: IEAKLTNTTTESRCPTQGEPSLNEEQDKRFICKHSMVDRGWGNGCGLFGKGGIVTCAMFT     120
DV2 THSTI : IEAKLTNTTTASRCPTQGEPSLNEEQDKRFVCKHSMVDRGWGNGCGLFGKGGIVTCAMFT     120
            ********** *******************:****************************

DV2 16681 : CKKNMEGKVVQPENLEYTIVITPHSGEEHAVGNDTGKHGKEIKITPQSSITEAELTGYGT     180
DV2 US-BID: CKKNMEGKVVLPENLEYTIVITPHSGEEHAVGNDTGKHGKEIKITPQSSITEAELTGYGT     180
DV2 THSTI : CKKNMEGKIVQPENLEYTIVITPHSGEENAVGNDTGKHGKEVKVTPQSSITEAELTGYGT     180
            ********:* *****************:************:*:****************

DV2 16681 : VTMECSPRTGLDFNEMVLLQMENKAWLVHRQWFLDLPLPWLPGADTQGSNWIQKETLVTF     240
DV2 US-BID: VTMECSPRTGLDFNEMVLLQMEDKAWLVHRQWFLDLPLPWLPGADTQGSNWIQKETLVTF     240
DV2 THSTI : VTMECSPRTGLDFNEMVLLQMENKAWLVHRQWFLDLPLPWLPGADTQGSNWIQKETLVTF     240
            **********************:*************************************

DV2 16681 : KNPHAKKQDVVVLGSQEGAMHTALTGATEIQMSSGNLLFTGHLKCRLRMDKLQLKGMSYS     300
DV2 US-BID: KNPHAKKQDVVVLGSQEGAMHTALTGATEIQMSSGNLLFTGHLKCRLRMDKLQLKGMSYS     300
DV2 THSTI : KNPHAKKQDVVVLGSQEGAMHTALTGATEIQMSSGNLLFTGHLKCRLRMDKLQLKGMSYS     300
            ************************************************************

DV2 16681 : MCTGKFKVVKEIAETQHGTIVIRVQYEGDGSPCKIPFEIMDLEKRHVLGRLITVNPIVTE     360
DV2 US-BID: MCTGKFKIVKEIAETQHGTIVIRVQYEGDGSPCKIPFEIMDLEKRHVLGRLITVNPIVTE     360
DV2 THSTI : MCTGKFKVVKEIAETQHGTIVIRVQYEGDGSPCKIPFEIMDLEKRHVLGRLITVNPIVTE     360
            *******:****************************************************

DV2 16681 : KDSPVNIEAEPPFGDSYIIIGVEPGQLKLNWFKKGSSIGQMFETTMRGAKRMAILGDTAW     420
DV2 US-BID: KDSPVNIEAEPPFGDSYIIIGVEPGQLKLNWFKKGSSIGQMFETTMRGAKRMAILGDTAW     420
DV2 THSTI : KDSPVNIEAEPPFGDSYIIIGVEPGQLKLSWFKKGSSIGQMFETTMRGAKRMAILGDTAW     420
            *****************************.******************************

DV2 16681 : DFGSLGGVFTSIGKALHQVFGAIYGAAFSGVSWTMKILIGVIITWIGMNSRSTSLSVTLV     480
DV2 US-BID: DFGSLGGVFTSIGKALHQVFGAIYGAAFSGVSWTMKILIGVIITWIGMNSRSTSLSVSLV     480
DV2 THSTI : DFGSLGGVFTSIGKALHQVFGAIYGAAFSGVSWTMKILIGVVITWIGMNSRSTSLSVSLV     480
            *****************************************:***************:**

DV2 16681 : LVGIVTLYLGVMVQA     495
DV2 US-BID: LVGVVTLYLGVMVQA     495
DV2 THSTI : LVGVVTLYLGVMVQA     495
            ***:***********
```

Conservative substitution

Non-conservative substitution

| % Identity/ % Similarity between E-protein sequence | DV2-16681 | DV2-US-BID | DV2-THSTI |
|---|---|---|---|
| DENV2-16881 | 100/100 | | |
| DV2-US/BID-V594/2006 | 98.384/99.59 | 100/100 | |
| DV2-THSTI/TRC/01 | 97.778/99.59 | 98/99 | 100/100 |

**Figure S1. Multiple sequence alignment between the E-proteins of DENV2 strains.** DV2 US-BID and DV2 THSTI refers respectively to DENV2-US/BID-V594/2006 and DENV2-

THSTI/TRC/01 strains which have been used in this study. DV2-16881 is a well-studied and frequently used DENV virus strain reference.

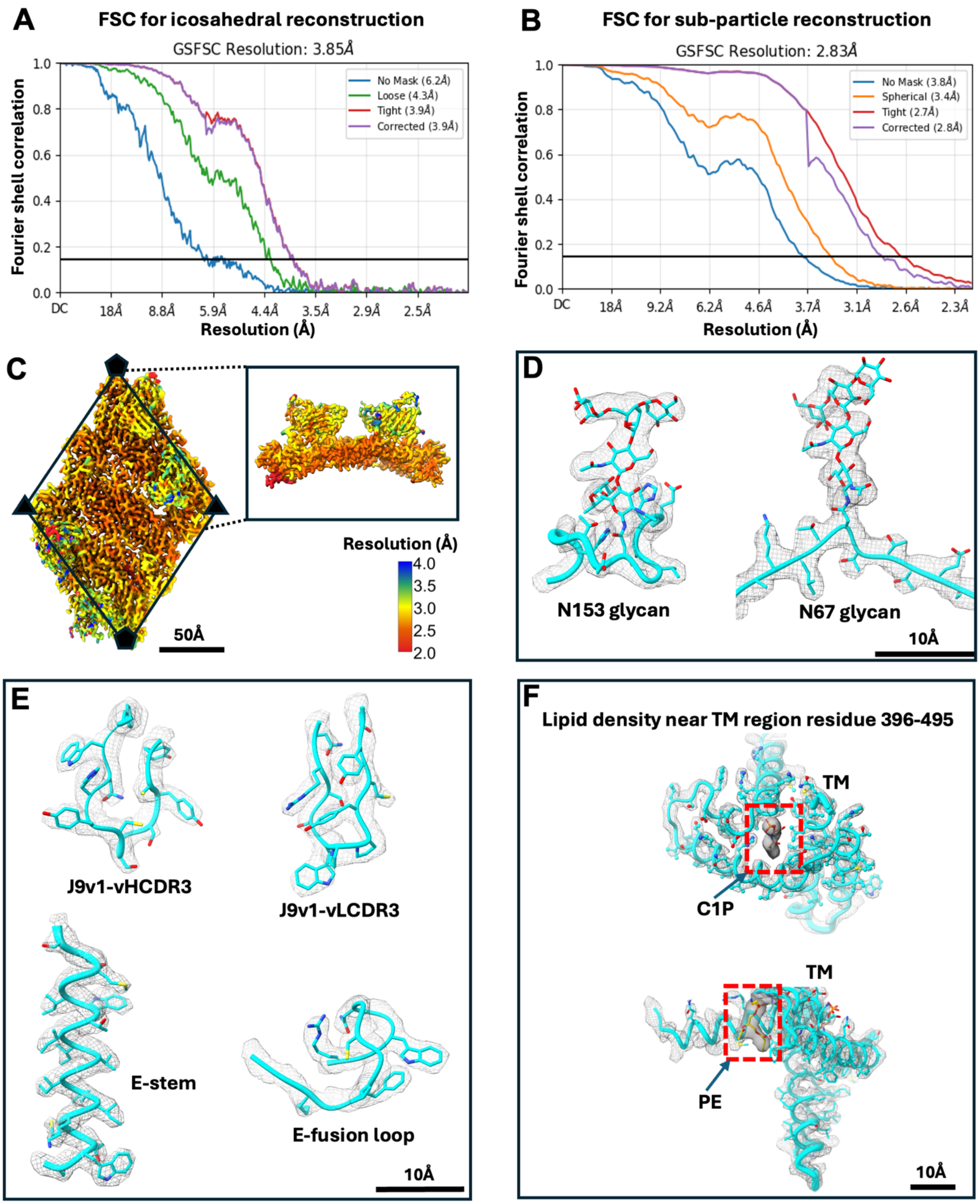

**Figure S2: Fourier shell correlation (FSC) and map features for cryo-EM reconstruction of J9 Fab complexed to DENV2-US/BID-V594/2006.** A. FSC of icosahedral reconstruction map.

B. FSC of sub-particle reconstruction. C. Local resolution estimate for the sub-particle reconstruction of E-dimer raft with Fab variable region. D-F. Map cutouts from the focused, sub-particle refined structure with built-in atomic models:- of the conserved glycans (D), different regions of DENV2 E-protein, variable heavy and light chain region of J9 (E) and lipid molecules ceraminde-1-phosphate (C1P) and PE(Phosphatidylethanolamine) density in the transmembrane region (TM) of the viral glycoproteins (F).

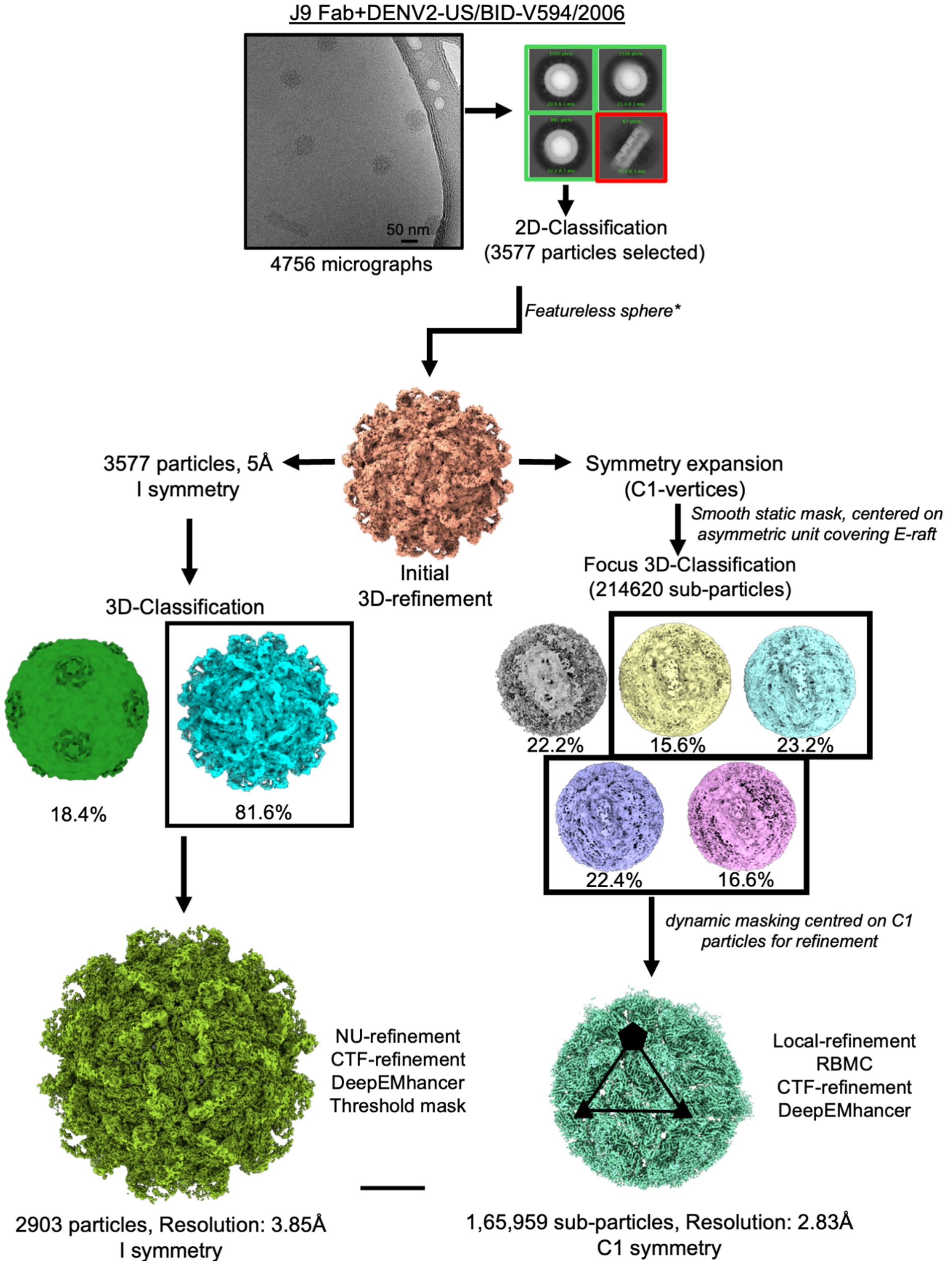

**Figure S3. Cryo-EM data processing workflow for J9 Fab complexed to icosahedral DENV2-US/BID-V594/2006.** *Featureless sphere was used as initial model in first round of 3D-refinement with icosahedral symmetry.

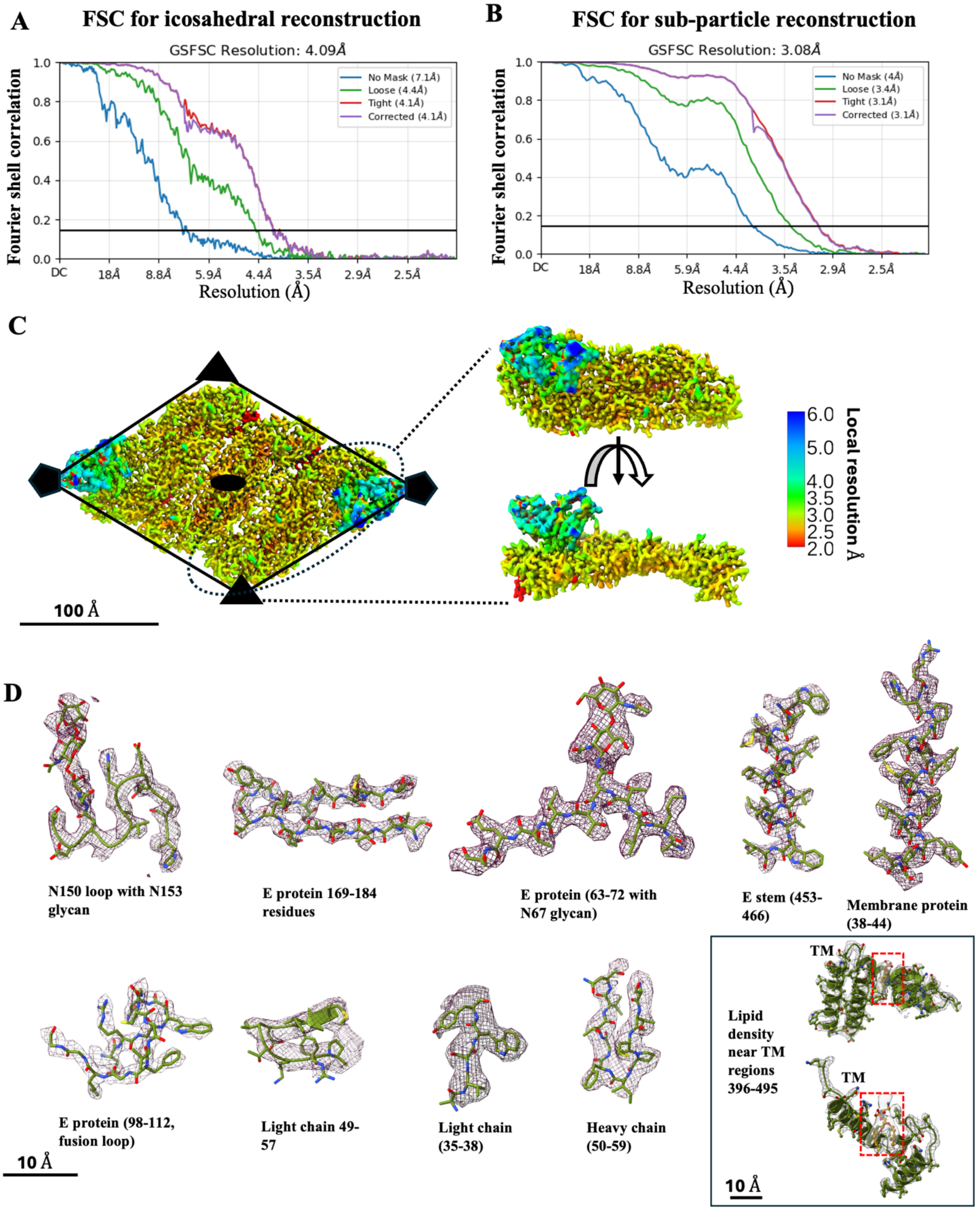

**Figure S4: Fourier shell correlation (FSC) and map features for cryo-EM reconstruction of D14.F25.S02 Fab complexed to DENV2-US/BID-V594/2006.** A. FSC of icosahedral reconstruction map. B. FSC of sub-particle reconstruction. C. Local resolution estimate for the sub-particle reconstruction of E-dimer raft with Fab variable region. D. Map cutouts from the focused, sub-particle refined structure with built-in atomic models from different regions of DENV2 E-protein and conserved glycans along with variable heavy and light chain region of Fab. The box shows docked lipid molecules in transmembrane region (TM) density of the viral glycoproteins.

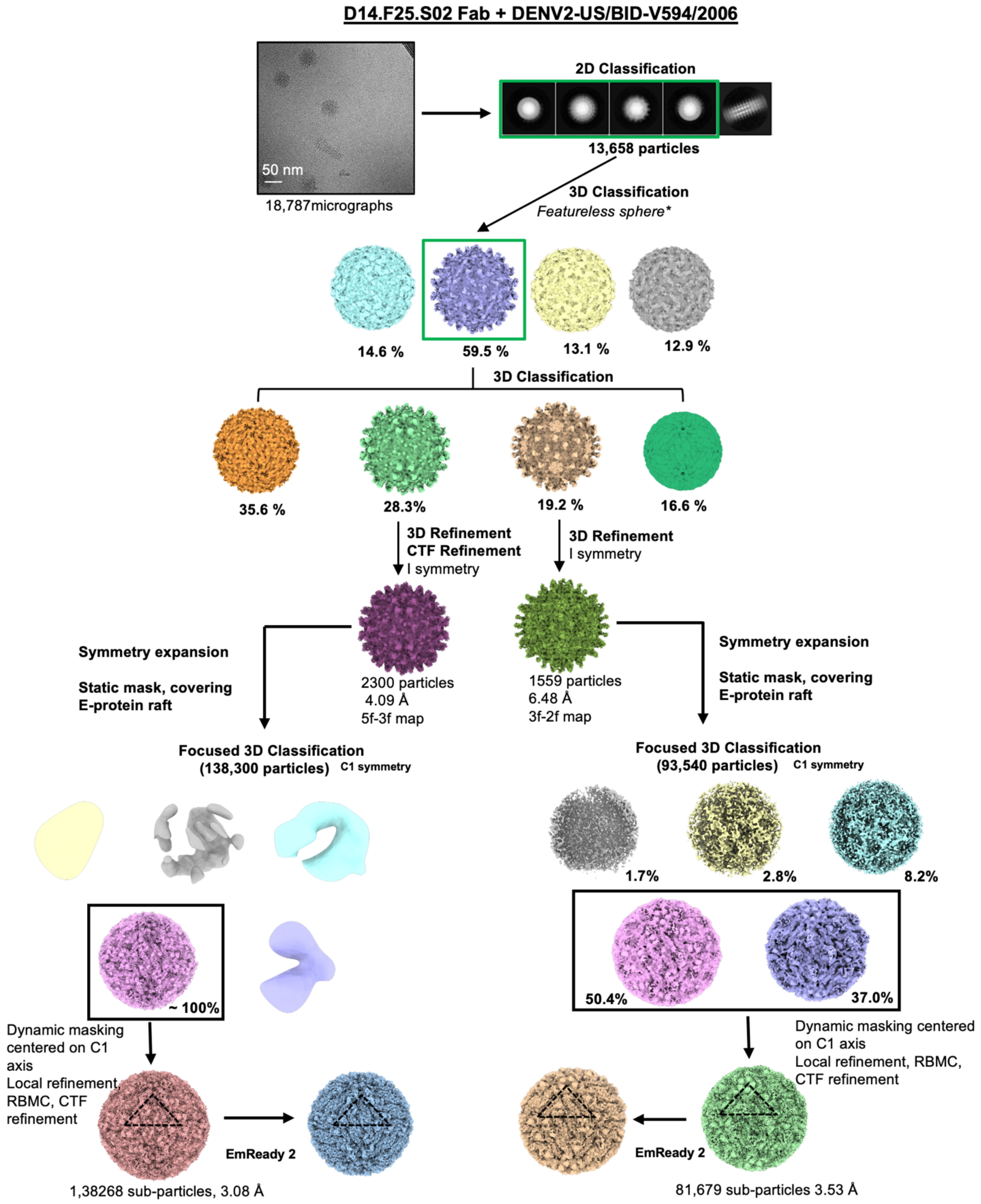

**Figure S5. Cryo-EM data processing workflow for D14.F25.S02 Fab complexed to icosahedral DENV2-US/BID-V594/2006.** *indicates initial model used.

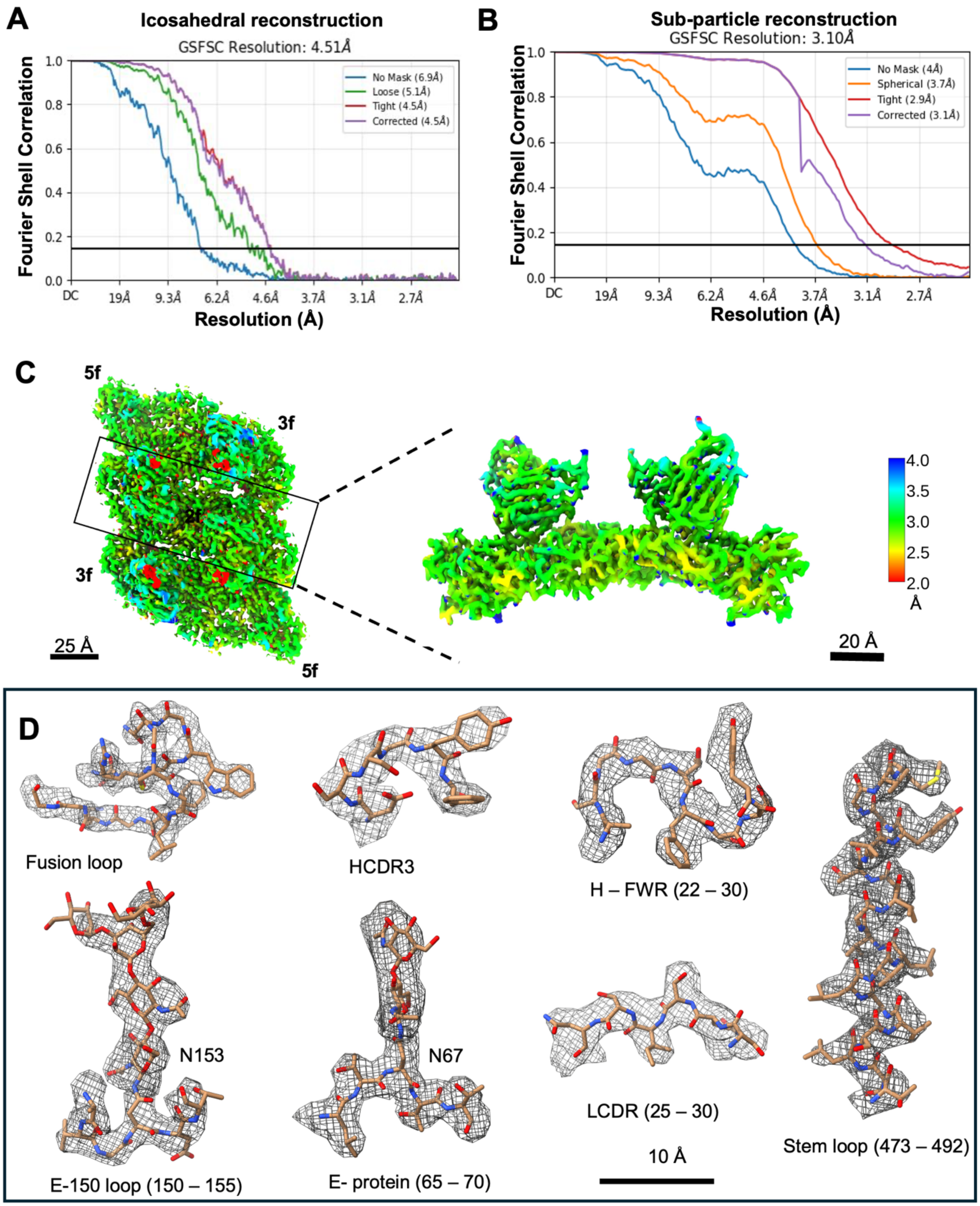

**Figure S6. Fourier shell correlation (FSC) and map features for cryo-EM reconstruction of D14.F05.S03 Fab complexed to DENV2-THSTI/TRC/01.** A and B. FSC curves for icosahedral and sub-particle reconstructions. C. Local resolution estimation of the sub-particle reconstruction of E-dimer raft with Fab variable region. D. Map cutouts from the focused, sub-particle refined structure with built-in atomic models from different regions of DENV2 E-protein and conserved glycans along with variable heavy and light chain regions of Fab.

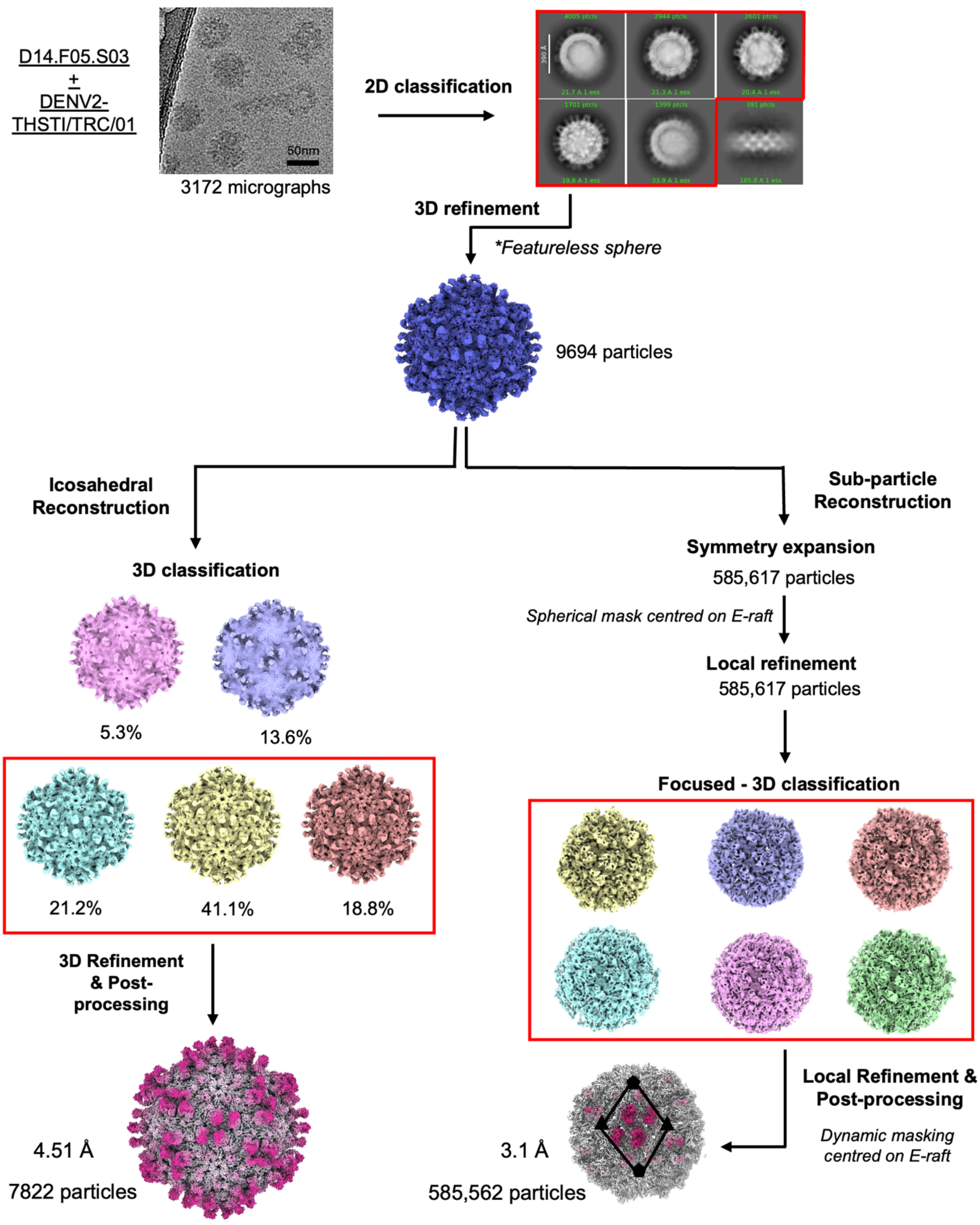

**Figure S7. Cryo-EM data processing workflow for D14.F05.S03 Fab complexed to icosahedral DENV2-THSTI/TRC/01.** * indicates the initial model used during 3D refinement.

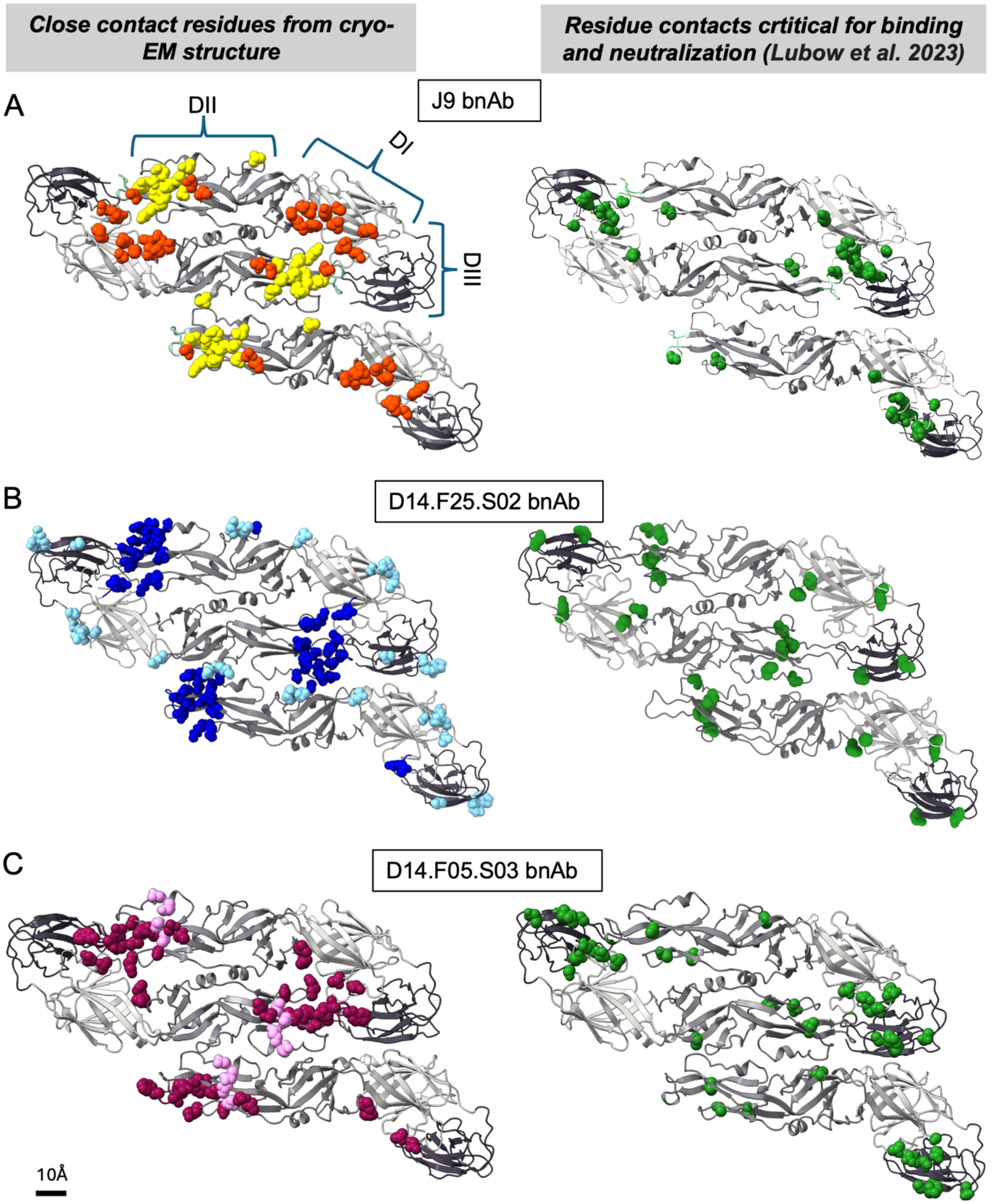

**Figure S8: HEDR bnAb contacts to the DENV E-protein. A-C**. Ribbon diagrams of E-proteins in an asymmetric unit of DENV are shown. Left: Solid balls indicate residues in close contact with bnAbs (inter-side-chain distance < 5Å). BnAb heavy and light chain contacts are respectively represented as orange and yellow in case of J9 bnAb (panel A), dark blue and light blue in case of D14.F25.S02 (panel B), dark pink and light pink in case of D14.F05.S03 (panel C). Right: Amino acid residues identified as important for neutralization and binding from previous functional analysis reports, coloured as green balls(*2*, *3*).

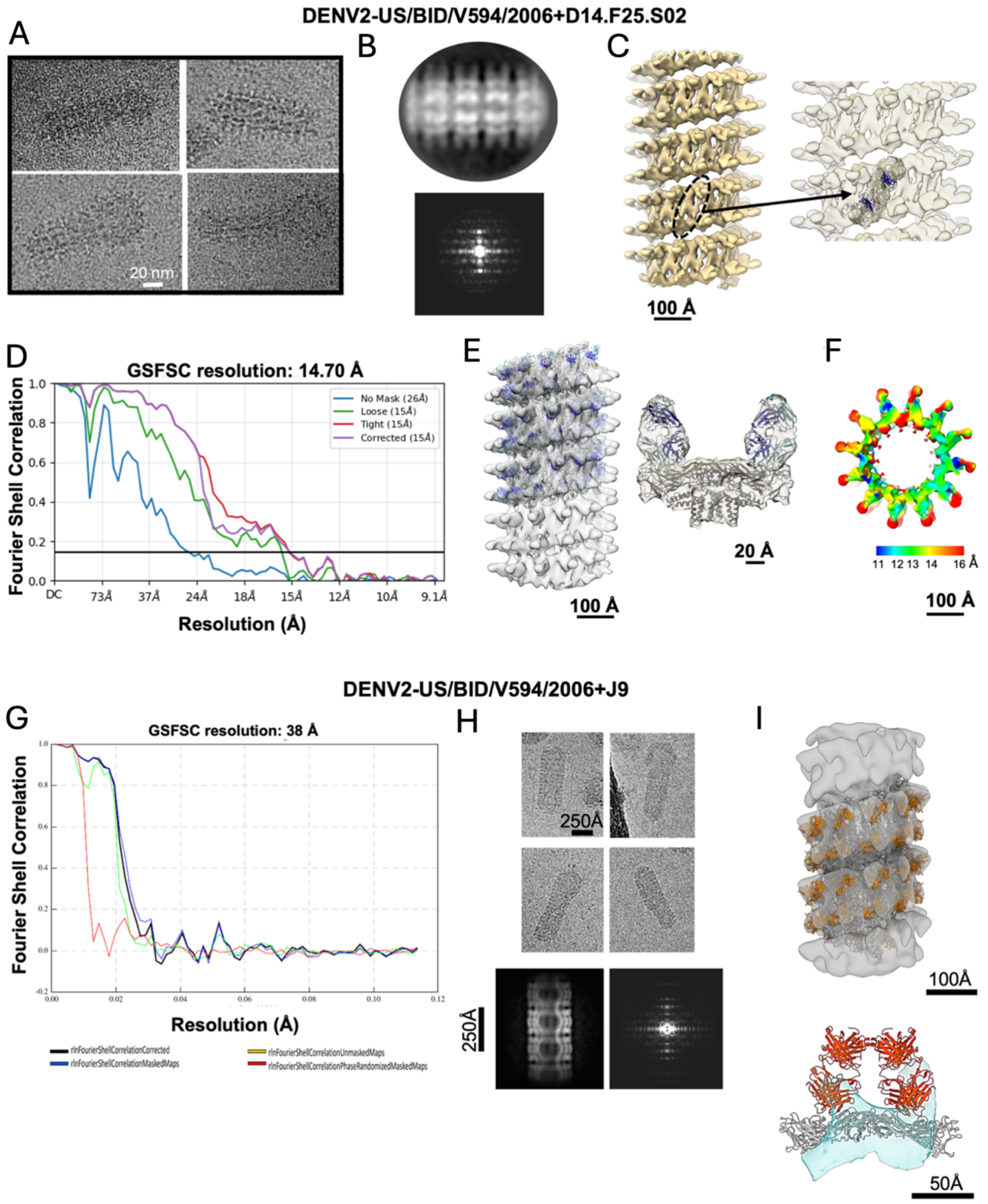

**Figure S9. Helical reconstructions of Fab-bound DENV2-US/BID-V594/2006 complexes.**

Panels A-F correspond to D14.F25.S02 Fab in complex with DENV2-US/BID-V594/2006.

A. Representative raw particle images showing DENV2-US/BID-V594/2006 particles which appear as complete tubes or with spherical heads and tubular tails.
B. Representative 2D class average showing helical repeats in the tubular virion regions along with power spectrum of the same which shows helical layer line pattern.
C. High threshold surface representation of helical cryo-EM reconstruction of D14.F25.S02-Fab bound virions (left), showing clear density repeats for the E-dimer and Fab. Black dotted oval indicates a single E-dimer with zoomed-in view (right) showing the fit of E-dimer atomic model into the density.
D. Fourier shell correlation curve for D14.F25.S02+DENV2-US/BID-V594/2006 3D reconstruction with imposed helical symmetry.
E. Helical map with fitted models of E-protein and Fab, along with individual cut-outs of one E-dimer bound to two Fabs.
F. Local resolution calculation.

Panels G-I correspond to J9 Fab in complex with DENV2-US/BID-V594/2006.

G. Fourier shell correlation curve for 3D reconstruction J9 Fab-bound tubular particles with imposed helical symmetry.
H. Representative raw images showing DENV2-US/BID-V594/2006 tubular particles along with a representative 2D class average showing helical repeats in the tubular virion regions and associated power spectrum of the same which shows helical layer line pattern.
I. Helical map with fitted models of E-protein and Fab, along with individual map cut-out fitted with one E-dimer and Fab variable region.

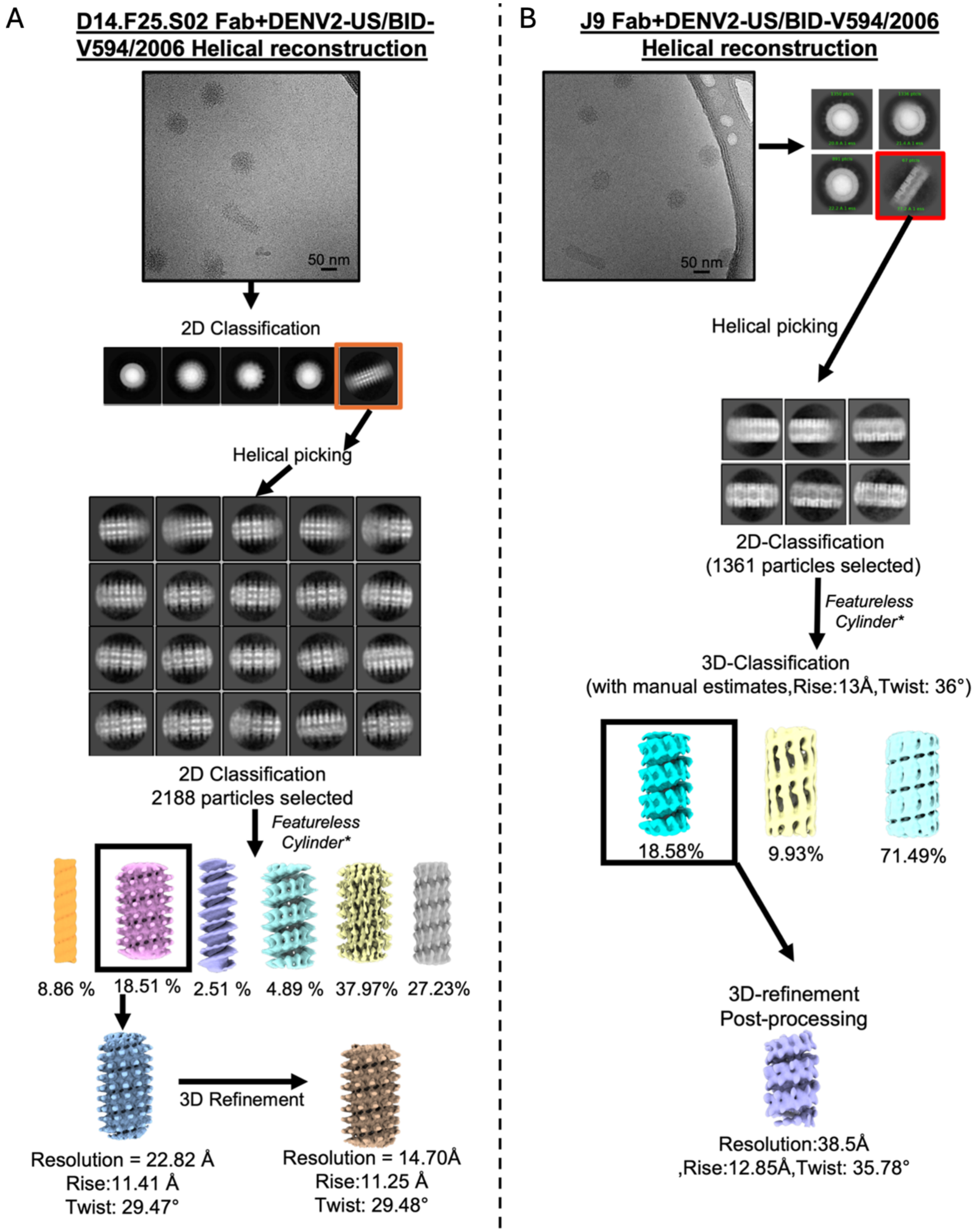

**Figure S10. Cryo-EM data processing workflow for Fab-bound tubular DENV2-US/BID-V594/2006 particles with helical symmetry (a and b).** *indicates the respective initial models used during 3D refinement and 3D-classification.

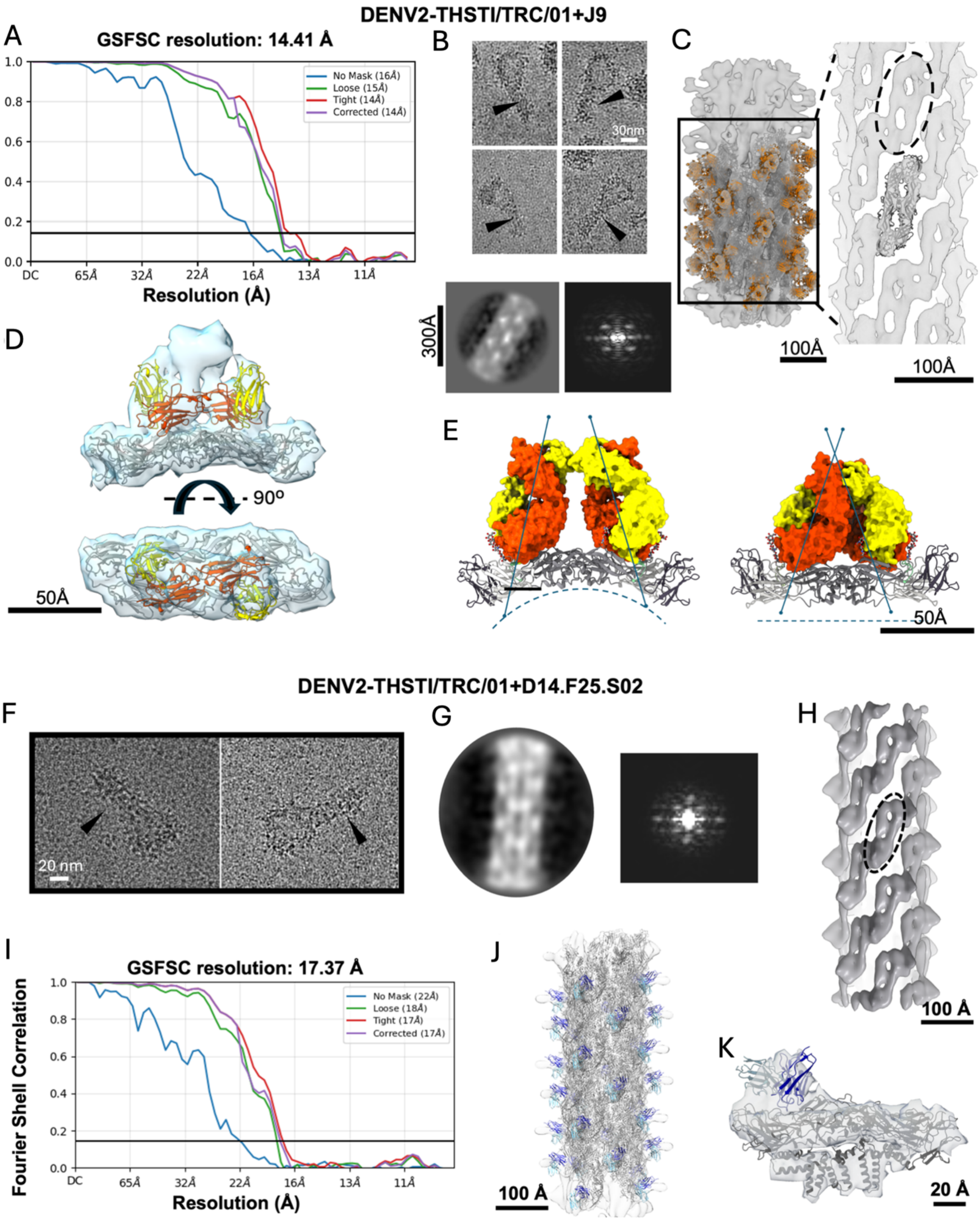

**Figure S11. Helical cryo-EM reconstructions of Fab-bound complexes with DENV2-THSTI/TRC/01.**

Panels A-E correspond to J9 Fab in complex with DENV2-THST/TRC/01.

A. Fourier shell correlation curve for helical structure of J9 Fab complexed with DENV2-THSTI/TRC/01 tubular particles.
B. Top: Representative images showing tubular J9-bound DENV2-THSTI/TRC/01 particles with black arrows indicating tubular regions. On micrograph top right shows the power spectrum which have similar pattern to helical layer lines. Bottom: Representative 2D class average with power spectrum showing layer lines.
C. Left: Rigid body fitting of E-protein and J9V Fab atomic models in helical map. Right: High threshold surface representation of the map showing clear density repeats for the E-dimer. Black dotted oval indicates a single E-dimer density with E-dimer atomic model fitted into the density below.
D. Density map cut-out showing good fit of E-dimer with both J9 Fab variable regions. Side-view shows clashing of Fab constant region densities in the EM map.
E. Left: Full J9 Fab positioned on icosahedral E-dimer (from high resolution cryo-EM model) does not show any clashes. Right: Full J9 Fab superimposed on its epitope in purified E-protein crystal structure (PDB: 6WY1) which has a flatter conformation of E-protein results in clashes of Fab constant domains as seen in the EM density maps in panels c,d.

Panels F-K correspond to D14.F25.S02 Fab in complex with DENV2-THST/TRC/01.

F. Representative raw particle images showing DENV2-THSTI/TRC/01 particles which appear as spherical heads with tubular tails.
G. Representative 2D class average showing helical repeats in the tubular virion regions along with power spectrum of the same which shows helical layer line pattern.

H. High threshold surface representation of helical cryo-EM reconstruction of D14.F25.S02-Fab bound virions showing clear density repeats for the E-dimer. Black dotted oval indicates a single E-dimer.

I. Fourier shell correlation curve for D14.F25.S02+DENV2-THSTI/TRC/01 3D reconstruction with imposed helical symmetry.

J. Helical map with fitted models of E-protein and Fab.

K. Density cut-out of one E-dimer bound to one Fab showing good fit of model to map.

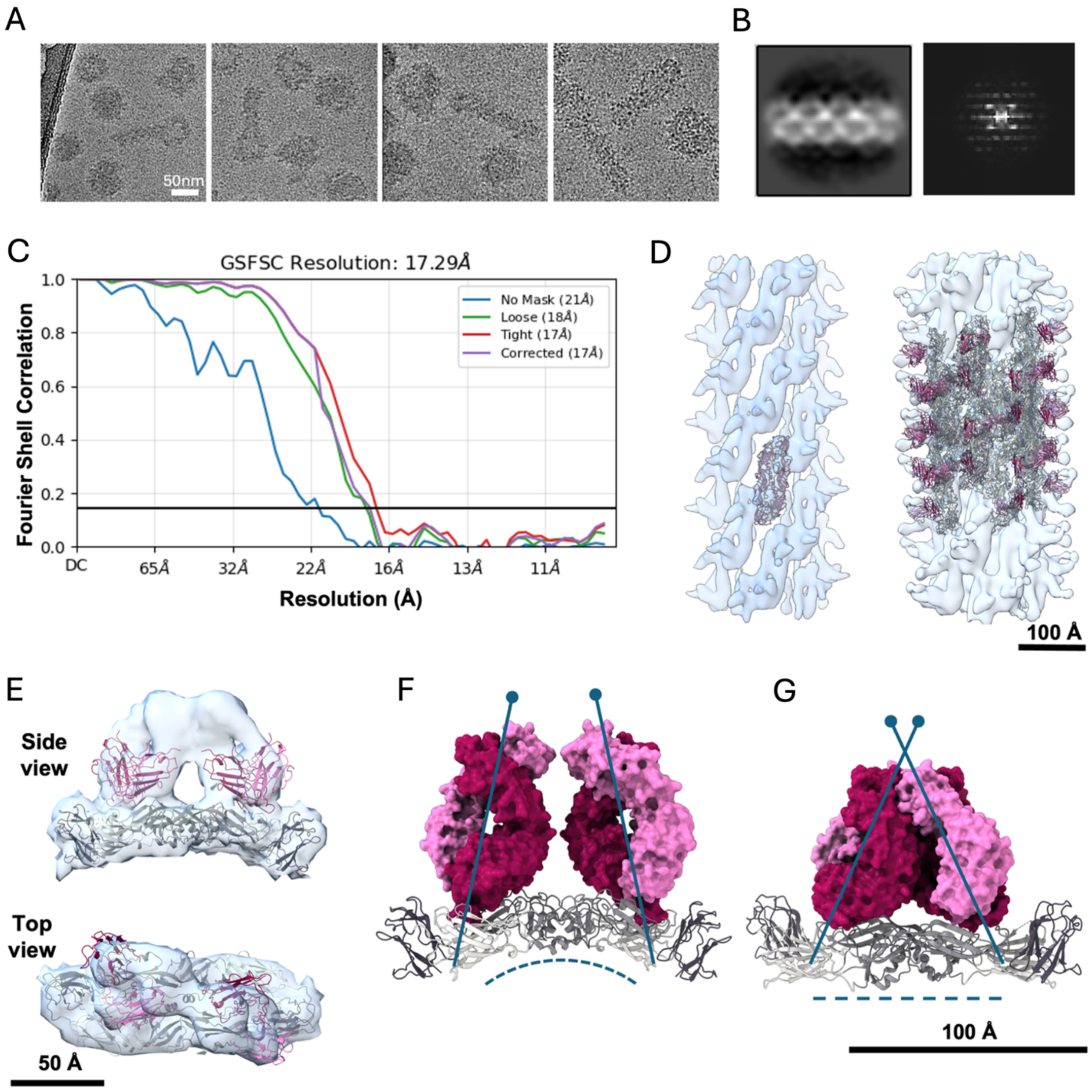

**Figure S12. Helical cryo-EM reconstructions of D14.F05.S03 in complex with DENV2-THSTI/TRC/01 strain.**

A. Representative micrographs showing DENV2-THSTI/TRC/01 virions with spherical head and tubular extensions. B. A representative 2D class and its power spectrum showing helical layer lines. C. Fourier shell correlation curve of helical reconstruction. D. Left: High threshold surface rendering showing clear densities for E-protein in reconstructed helical map with one E-dimer fitted for reference. Right: Rigid body fitting of E-proteins and Fab models into the helical map. E. Map density cutout showing fitted E-dimer with Fab variable chain model. F. Fab arrangement in icosahedral E-dimer structure where no clash is observed between Fab constant domains. G. Fab superimposed on its epitope in purified E-protein crystal structure (PDB: 6WY1) which has a flatter conformation of E-protein. In this flat E-dimer conformation, binding of Fabs leads to clashes in Fab constant domains.

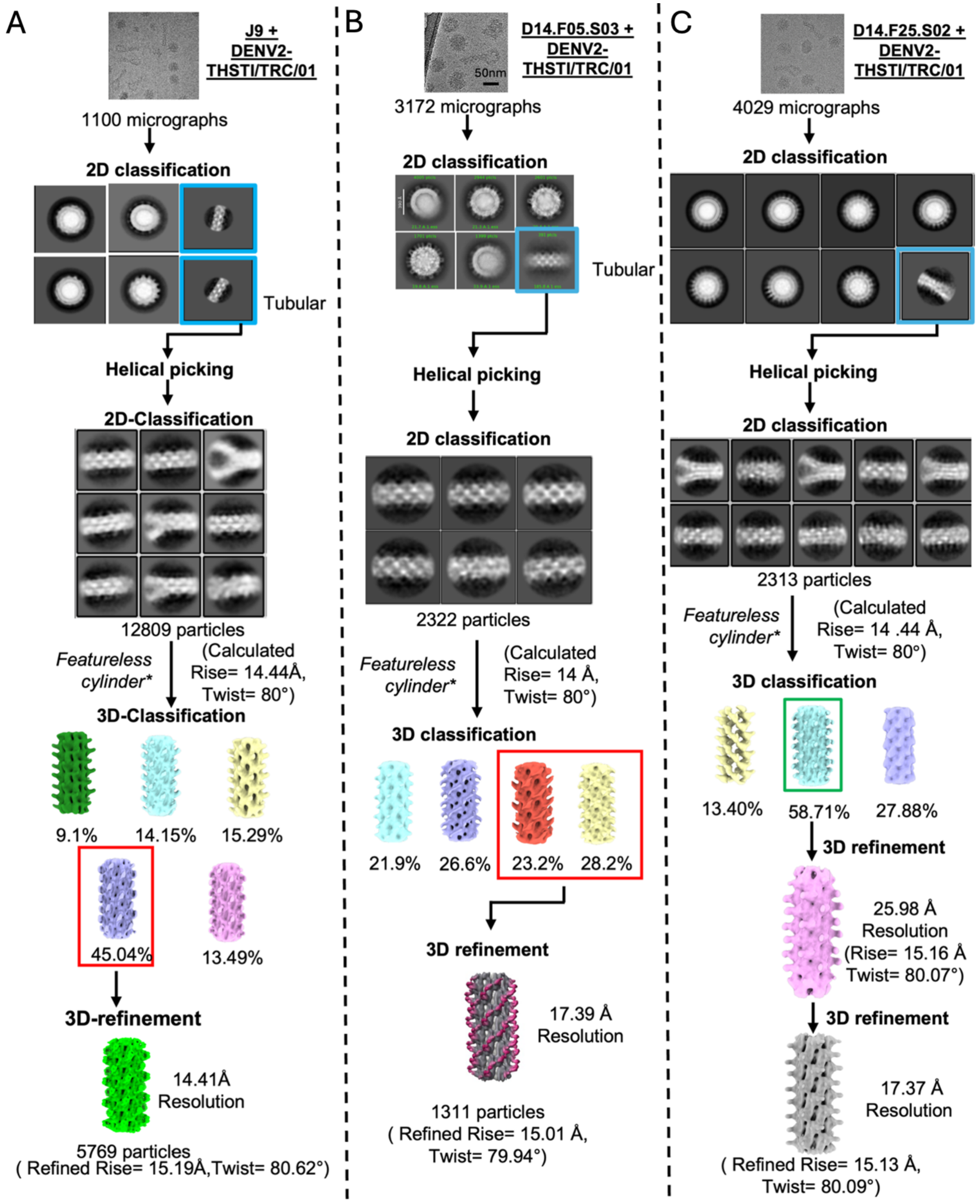

**Figure S13. Cryo-EM data processing workflow for structures of Fab-bound tubular DENV2-THSTI/TRC/01 with helical symmetry (A-C).** *indicates respective initial models

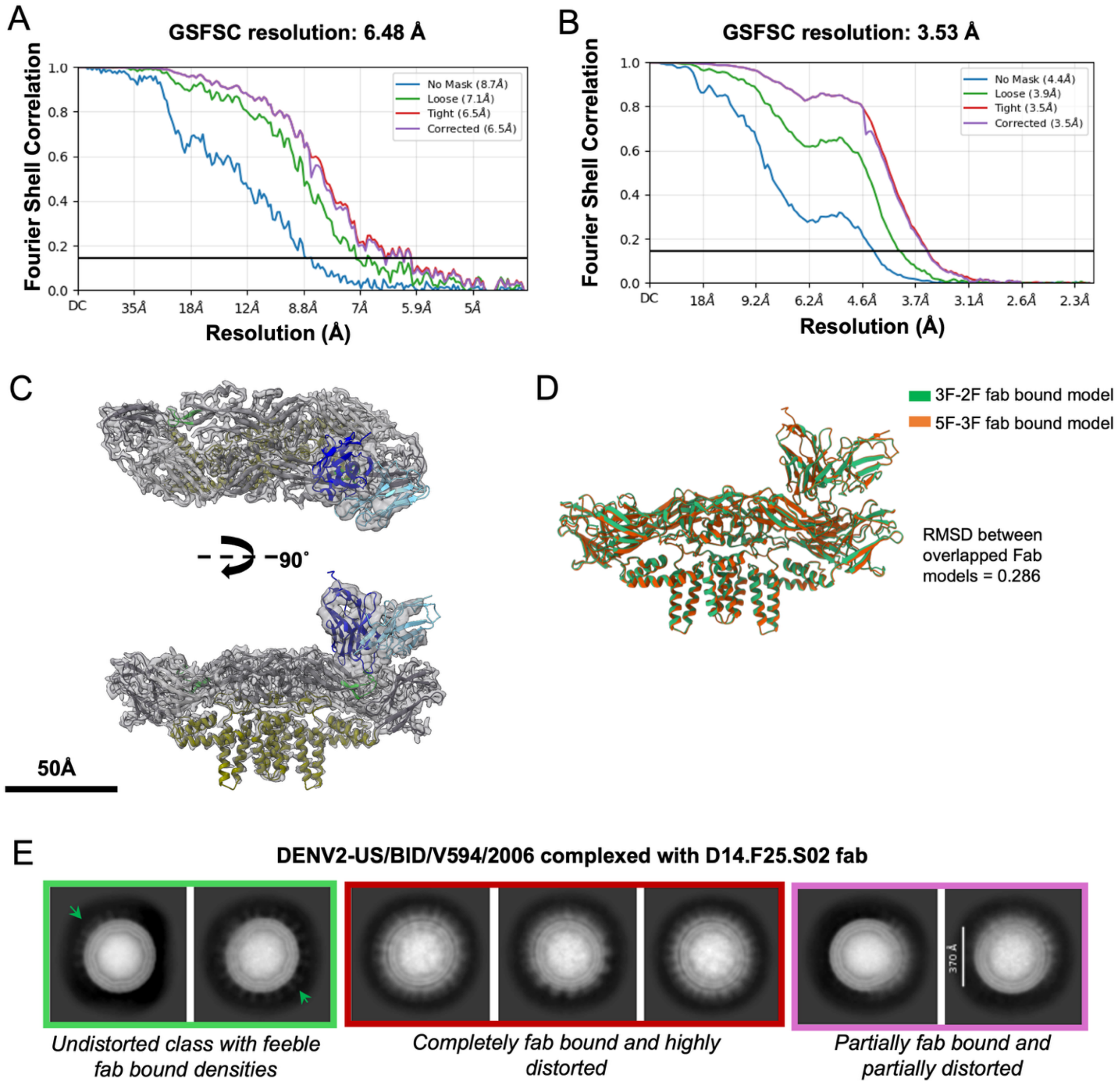

**Figure S14. Cryo-EM reconstruction and analyses of 3f-2f map of DENV2-US/BID/V594/2006 complexed to D14.F25.S02 Fab.** A. Fourier shell correlation curve plot of 2f-3f icosahedral reconstruction. B. Fourier shell correlation curve of sub-particle refinement. C. Density cut-out of an asymmetric unit showing built-in atomic models E-protein and Fab. D. Overlap of the atomic models of E-dimer+Fab from the 3f-2f map and 5f-3f map showing nearly identical structures of the Fab and complex. E. 2D class averages of DENV2-US/BID-

V594/2006 complexed with D14.F25.S02 Fab. Only classes with overall mature appearance are shown. Classes are labelled with green, pink and red colored square based on extent of virion distortion.

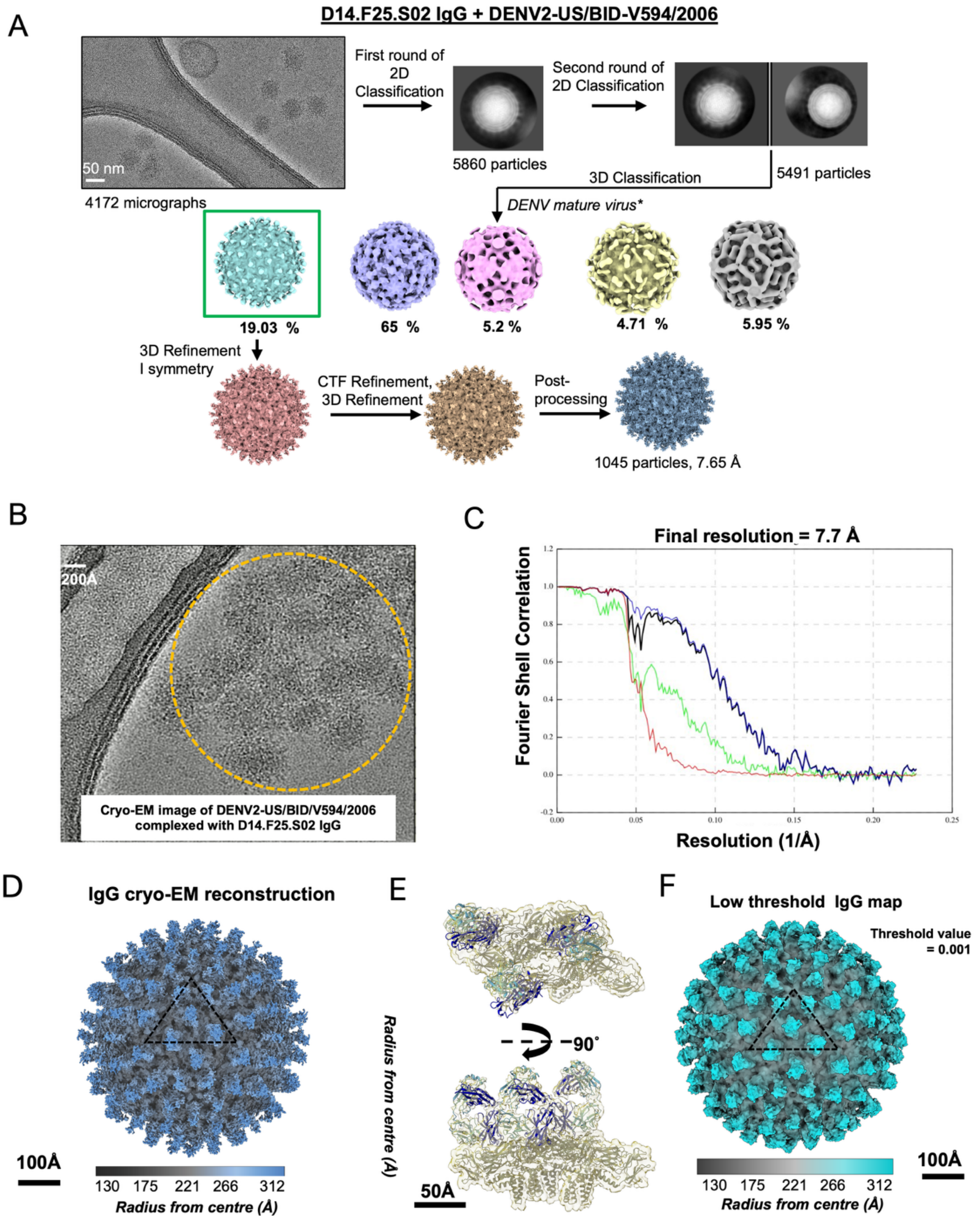

**Figure S15. Cryo-EM reconstruction and analyses of D14.F25.S02 IgG bound to DENV2-US/BID/V594/2006.** A. Cryo-EM data processing workflow for D14.F25.S02 IgG bound to spherical DENV. *indicates initial model used during 3D classification. B. Representative raw cryo-EM micrograph of DENV2-US/BID-V594/2006 complexed with D14.F25.S02 IgG showing aggregation of virus particles. C. Fourier shell correlation of icosahedral reconstruction IgG complex. D. Cryo-EM reconstruction of IgG complex. E. Density cut-out of an asymmetric unit showing good fit of E-dimers and Fab models. F. Very low threshold IgG map to show the absence of density for the Fc regions of IgG.

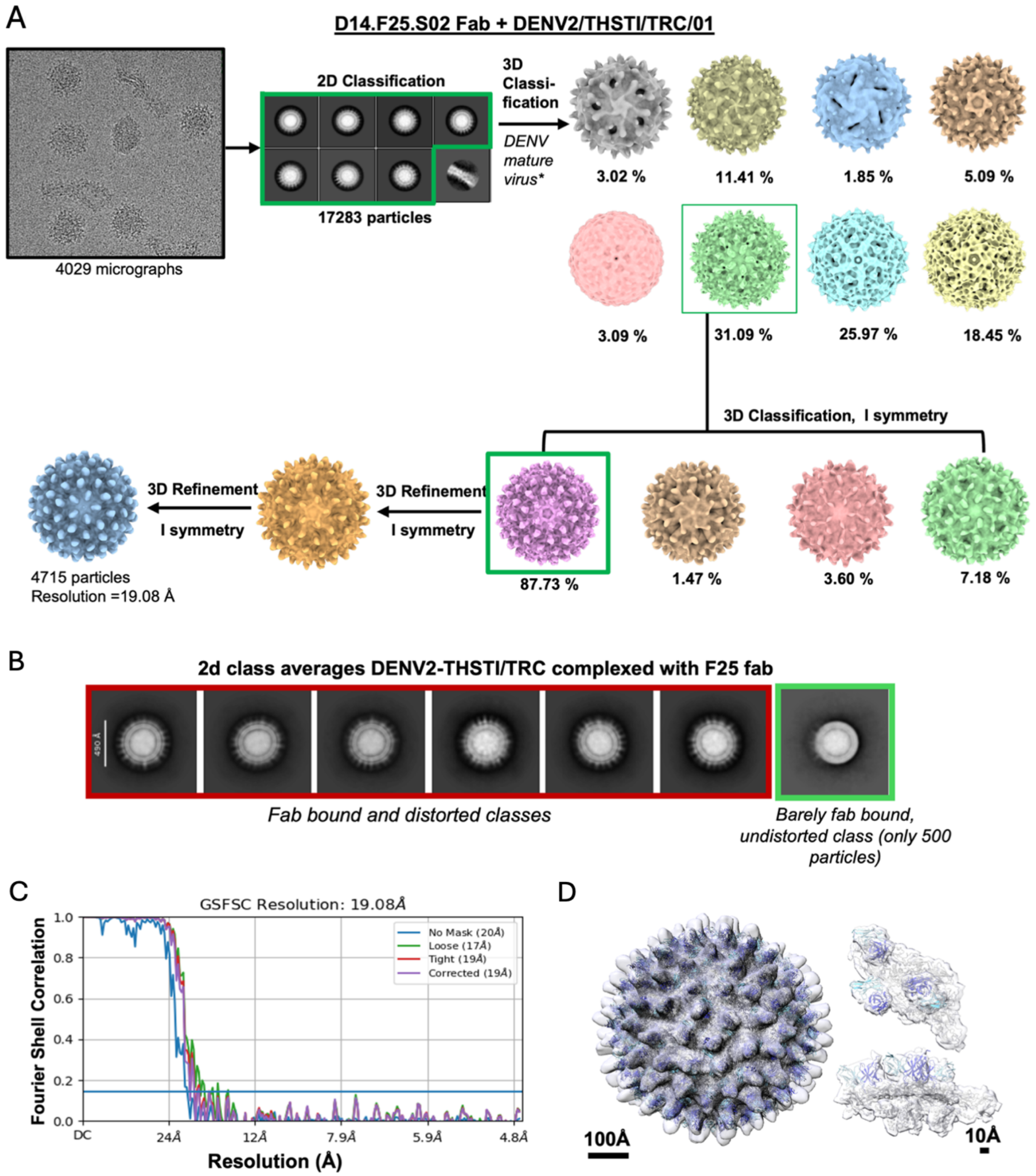

**Figure S16. Cryo-EM structure analysis of D14.F25.S02 complexed to DENV2-THSTI/TRC/01 strain.** A. Cryo-EM data processing workflow for D14.F25.S02 Fab bound to spherical DENV2-THSTI/TRC/01 strain. *indicates initial model used during 3D classification.

B. 2D class averages of spherical DENV2-THSTI/TRC/01 virions complexed with D14.F25.S02 Fab. Only classes with overall mature appearance are shown. Distorted and undistorted classes have been labelled by red and green squares respectively. C. Fourier shell correlation plot of icosahedral reconstruction of D14.F25.S03 + DENV2-THSTI/TRC/01 strain. D. Cryo-EM density map of D14.F25.S03 Fab + DENV2-THSTI/TRC/01 with atomic models of E-proteins and D14.F25.S02 Fab fitted into it as rigid bodies. Map cutouts of asymmetric unit are shown to highlight the good fitting of model to map.

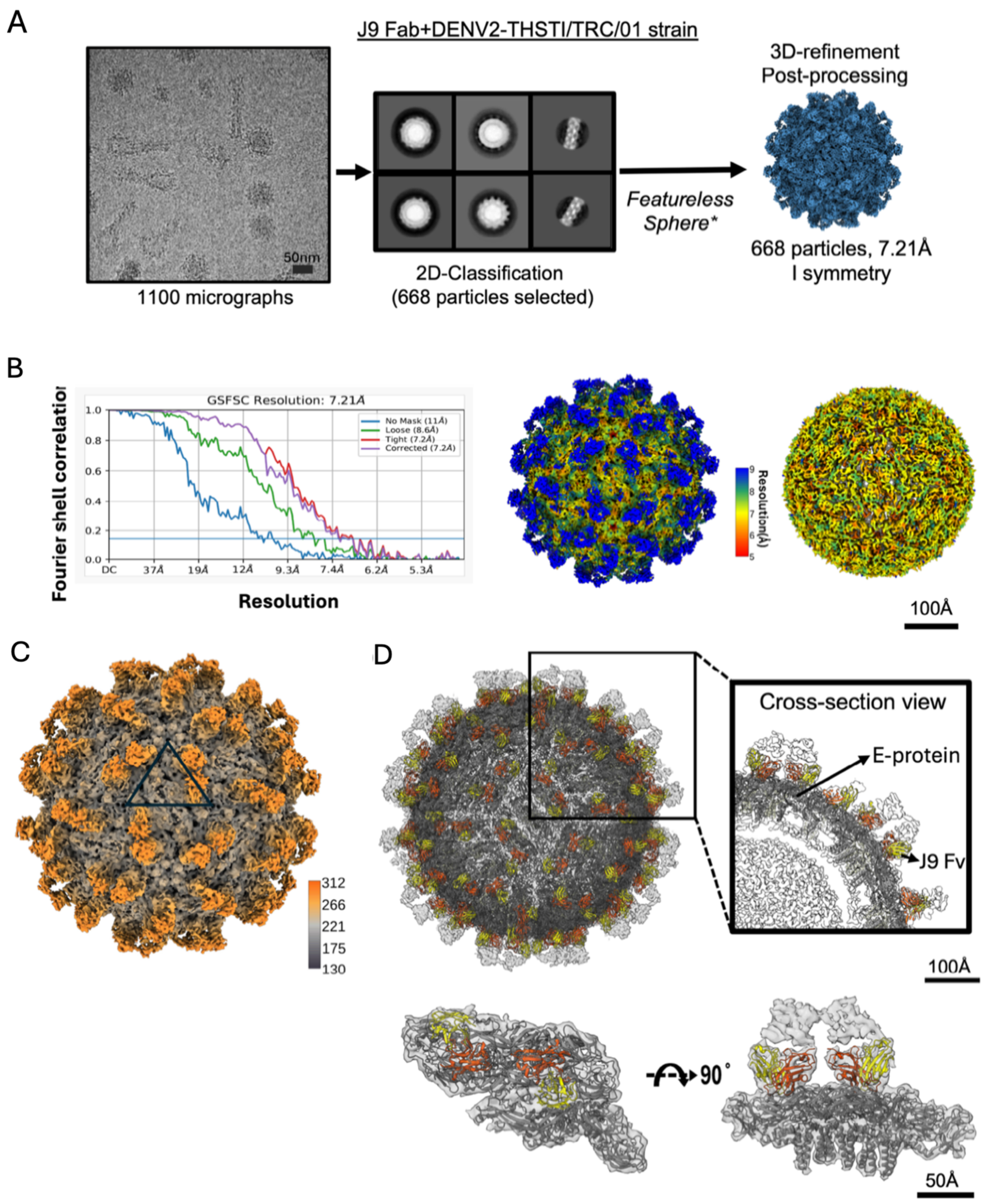

**Figure S17. Cryo-EM reconstruction and map details of J9 Fab + spherical DENV2-THSTI/TRC/01 strain.** A. Cryo-EM data processing workflow for J9 Fab bound to spherical DENV2-THSTI/TRC/01 strain. *indicates initial model used during 3D classification.

B. Left: Fourier shell correlation curve for icosahedral reconstruction of J9 fab complex. Right: Local resolution estimation of the cryo-EM map. C. Surface representation of cryo-EM map with radial coloring. Icosahedral asymmetric unit is denoted as a black triangle. D. Rigid body fitting of E-protein (grey ribbons) and J9 fab variable regions (orange and yellow ribbons representing the heavy and light chain of the Fab respectively) into the icosahedral map. Zoom in showing map to model fit for a cross-section of virus (top right). Fitting of E-asymmetric unit and J9 fab region map-zoned from the full icosahedral structure of virus and fab bound complex (bottom).

**Table S1.** Cryo-EM data collection, refinement and validation statistics for DENV2 complexed with J9 bnAb.

a. J9 complexes with DENV2-US/BID-V594/2006 strain.

| | **DENV2/US/HBID-V594/2006+J9 fab complex** | | |
|---|---|---|---|
| **Data collection and processing** | | | |
| Magnification | 81,000 | | |
| Voltage (kV) | 300 | | |
| Electron exposure (e–/Å$^2$) | 40.3 | | |
| Defocus range (μm) | -1.0 to -2.5 | | |
| Pixel size (Å) | 1.1 | | |
| | **EMDB: 63759 PDB: 9MAV** | **EMDB: 80279 PDB: 25OV** | **EMDB: 63762 PDB: 9MAZ** |
| Mode of Reconstruction | Icosahedral reconstruction | Sub-particle reconstruction | Helical Reconstruction |
| Symmetry imposed | Icosahedral symmetry (I1) | C1 symmetry | Helical (Point group C1) Rise=12.83Å Twist=35.83° |
| Initial particle images (no.) | 15000 | 214620 (sub-particles) | 2450 (segments) |
| Final particle images (no.) | 2903 | 165959 (sub-particles) | 253 (segments) |
| FSC threshold | 0.143 | 0.143 | 0.143 |
| Map resolution (Å) | 3.85 | 2.83 | 38.5 |
| Map resolution range (Å) | 2.5-6.5 | 2.2-4.0 | N/A |
| **Model Refinement and validation** | | | |
| Initial model used (PDB code) | High-resolution model (EMProt) | De-novo building (EMProt) | Atomic model from high resolution icosahedral structure. |
| Model resolution (Å)* | 3.3 | 1.8 | N/A |
| D Model* | 3.8 | 2.3 | N/A |
| Map sharpening *B* factor (Å$^2$) | N/A | N/A | N/A |
| Model composition | | | |
| Non-hydrogen atoms | 16772 | 25902 | |
| Protein residues | 2152 | 1602 | 1227 |
| Ligands | NAG:11 BMA:2 MAN:2 | NAG:8 BMA:4 FUC:2 MAN:8 | |

| | | | |
|---|---|---|---|
| R.m.s. deviations | | | N/A |
| Bond lengths (Å) | 0.003 | 0.004 | |
| Bond angles (°) | 0.759 | 0.625 | |
| Validation | | | N/A |
| MolProbity score | 1.78 | 1.35 | |
| Clashscore | 10.45 | 2.87 | |
| Poor rotamers (%) | 1.71 | 1.41 | |
| Ramachandran plot | | | N/A |
| Favored (%) | 97.7 | 97.1 | |
| Allowed (%) | 2.3 | 2.9 | |
| Disallowed (%) | 0 | 0 | |

b. J9 complexes with DENV2-THSTI/TRC/01 strain.

| | **DENV2/THSTI/TRC/01+J9 fab complex** | |
|---|---|---|
| **Data collection and processing** | | |
| Magnification | 36000 | |
| Voltage (kV) | 200 | |
| Electron exposure (e–/Å$^2$) | 48 | |
| Defocus range (μm) | -1.25 to -2.75 | |
| Pixel size (Å) | 1.16 | |
| | **EMDB: 63763**<br>**PDB: 9MB0** | **EMDB: 63764**<br>**PDB: 9MB1** |
| Mode of Reconstruction | Icosahedral reconstruction | Helical Reconstruction |
| Symmetry imposed | Icosahedral symmetry (I1) | Helical (Point group C1)<br>Rise=15.19Å<br>Twist=80.62° |
| Initial particle images (no.) | 10108 | 19729 (segments) |
| Final particle images (no.) | 668 | 5769 (segments) |
| FSC threshold | 0.143 | 0.143 |
| Map resolution (Å) | 7.21 | 14.41 |

| | | |
|---|---|---|
| Map resolution range (Å) | N/A | N/A |
| **Model Refinement and validation** | Rigid Body Fitting | |
| Initial model used (PDB code) | Atomic model from high resolution icosahedral structure. | Atomic model from high resolution icosahedral structure. |
| Map sharpening *B* factor ($Å^2$) | -511.91 | N/A |
| Model composition | | |
| Protein residues | 2375 | 1227 |

**Table S2.** Cryo-EM data collection, refinement and validation statistics for DENV2 complexed with D14.F25.S02 bnAb.

a. D14.F25.S02 complexes with DENV2-US/BID-V594/2006 strain.

| | DENV2-US/BID-V594/2006 + D14.F25.S02 Fab | | | | | DENV2-US/BID-V594/2006+ D14.F25.S02 IgG |
|---|---|---|---|---|---|---|
| **Data collection and processing** | | | | | | |
| Magnification | 81,000 | | | | | 81,000 |
| Voltage (kV) | 300 | | | | | 300 |
| Electron exposure (e–/Å$^2$) | 40 | | | | | 40 |
| Defocus range (µm) | -1.0 to -2.5 | | | | | -1.0 to -2.5 |
| Pixel size (Å) | 1.1 | | | | | 1.1 |
| | **EMDB: 63643 PDB: 9M5H** | **EMDB: 63813 PDB: 9U38** | **EMDB:80390 PDB:25US** | **EMDB:80426 PDB:25WK** | **EMDB: 63809 PDB: 9U34** | **EMDB: 63812 PDB: 9U37** |
| Mode of Reconstruction | **Icosahedral reconstruction** | | **Sub-particle reconstruction** | | **Helical Reconstruction** | **Icosahedral reconstruction** |
| Symmetry imposed | Icosahedral symmetry (I1) 5f-3f map | Icosahedral symmetry (I1) 3f-2f map | C1 Symmetry | | Helical (Point group C1) Rise=11.25 Å Twist= 29.48° Order = 12 | Icosahedral symmetry (I1) |
| | | | 5f-3f map | 3f-2f map | | |
| Initial particle images (no.) | 46826 | 46826 | 138300 (sub-particles) | 93540 (sub-particles) | 4784 (segments) | 8151 |
| Final particle images (no.) | 2305 | 1559 | 138268 (sub-particles) | 81679 (sub-particles) | 405 (segments) | 1045 |
| FSC threshold | 0.143 | 0.143 | 0.143 | 0.143 | 0.143 | 0.143 |
| Map resolution (Å) | 4.09 | 6.48 | 3.08 | 3.53 | 14.7 | 7.65 |
| Map resolution range (Å) | 2.5-7.5 | 4.5-10.5 | 2.3-6 | 2.3-7.3 | 11-16 | 4 -10 |

| **Model Refinement and validation** | | | | | | |
|---|---|---|---|---|---|---|
| Initial model used (PDB code) | Atomic model from high resolution sub-particle reconstruction | Atomic model from high resolution sub-particle reconstruction | De-novo building (EMProt for Viral protein) and AlphaFold3 for Fab | De-novo building (EMProt for Viral protein) and AlphaFold3 for Fab | Atomic model from high resolution sub-particle reconstruction | Atomic model from high resolution sub-particle reconstruction |
| Model resolution (Å)* | N/A | N/A | 2.3 | 2.5 | N/A | N/A |
| FSC threshold | N/A | N/A | 0.143 | 0.143 | N/A | N/A |
| **Model composition** | | | | | | |
| Non-hydrogen atoms | N/A | N/A | 21400 | 21420 | N/A | N/A |
| Protein residues | N/A | N/A | 1379 | 1379 | N/A | N/A |
| Ligands | N/A | N/A | NAG:7<br>BMA:1 | NAG:7<br>BMA:1 | N/A | N/A |

b. D14.F25.S02 complexes with DENV2-THSTI/TRC/01 strain.

| **Data collection and processing** | **DENV2/THSTI/TRC/01 +D14.F25.S02 Fab complex** | |
|---|---|---|
| Magnification | 36000 | |
| Voltage (kV) | 200 | |
| Electron exposure (e–/Å$^2$) | 48 | |
| Defocus range (μm) | -1.25 to -2.75 | |
| Pixel size (Å) | 1.16 | |
| | **EMDB: 63811 PDB: 9U36** | **EMDB: 63810 PDB: 9U35** |
| Mode of Reconstruction | Icosahedral reconstruction | Helical Reconstruction |
| Symmetry imposed | Icosahedral symmetry (I1) | Helical (Point group C1) Rise=15.13 Å Twist=80.09° Order =9 |
| Initial particle images (no.) | 26782 | 8264 (segments) |
| Final particle images (no.) | 4715 | 1358 (segments) |

| | | |
|---|---|---|
| FSC threshold | 0.143 | 0.143 |
| Map resolution (Å) | 19.08 | 17.37 |
| Model used for rigid body fitting (PDB code) | Atomic model from high resolution sub-particle reconstruction | Atomic model from high resolution sub-particle reconstruction |

**Table S3**. Cryo-EM data collection, refinement and validation statistics for DENV2 complexed with D14.F05.S03 bnAb.

| | **DENV2/THSTI/TRC/01 + D14.F05.S03 Fab complex** | | |
|---|---|---|---|
| **Data collection and processing** | | | |
| Magnification | 36000 | | |
| Voltage (kV) | 200 | | |
| Electron exposure (e–/Å$^2$) | 38 | | |
| Defocus range (μm) | -1.25 to -2.5 | | |
| Pixel size (Å) | 1.16 | | |
| | **EMDB: 80506<br>PDB: 26BC** | **EMDB: 80540<br>PDB: 26CL** | **EMDB: 80544<br>PDB: 26CR** |
| **Mode of reconstruction** | **Icosahedral** | **Sub-particle** | **Helical** |
| Symmetry imposed | Icosahedral symmetry (I1) | C1 | Helical (C1 – point group)<br>Rise – 15.01 Å<br>Twist – 79.94°<br>Order - 9 |
| Initial particle images (no.) | 31068 | 5,81,639 | 8045 |
| Final particle images (no.) | 7822 | 5,81,636 | 1311 |
| Map resolution (Å)<br>FSC threshold: 0.143 | 4.51 | 3.1 | 17.29 |
| Map resolution range (Å) | 3-7 | 2.5-5 | N/A |
| **Refinement** | | | |
| Initial model used (PDB code) | high-resolution atomic model from sub-particle reconstruction | Denovo building (EMProt) | high-resolution atomic model from sub-particle reconstruction |
| Model resolution (Å)<br>FSC threshold: 0.143 | 4.4 | 2.1 | N/A |
| **Model composition** | | | |
| Non-hydrogen atoms | 15018 | 37467 | N/A |
| Protein residues | 1942 | 2406 | N/A |
| Ligands | NAG – 6 | NAG – 12; BMA – 3 ; MAN - 6 | N/A |
| **R.m.s. deviations** | | | |
| Bond lengths (Å) | 0.003 | 0.004 | N/A |
| Bond angles (°) | 0.644 | 0.679 | N/A |
| **Validation** | | | |
| MolProbity score | 2.02 | 1.80 | N/A |
| Clashscore | 12.00 | 5.47 | N/A |
| Poor rotamers (%) | N/A | N/A | N/A |
| **Ramachandran plot** | | | |
| Favored (%) | 93.51 | 95.17 | N/A |
| Allowed (%) | 6.49 | 4.83 | N/A |
| Disallowed (%) | 0.00 | 0.00 | N/A |

**Table S4.** Close contacts between J9 bnAb and DENV E-proteins based on high resolution cryo-EM maps and atomic model.

| **J9 Fv Heavy chain residues** | **E-dimer residues (E-E')** | **Distance ( all atom cutoff <5Å)** |
|---|---|---|
| Vh ALA 104 | E' ASN 103 | 3.032 |
| Vh ASP 63 | E' THR 66 | 3.02 |
| Vh SER 105 | E' GLY 102 | 4.223 |
| Vh THR 58 | E' LYS 247 | 4.93 |
| Vh SER 103 | E ASP 154 | 2.916 |
| Vh ASN 30 | E GLY 156 | 4.157 |
| Vh SER 31 | E THR 155 | 2.738 |
| Vh GLY 57 | E SER 273 | 3.965 |
| Vh PHE 55 | E LYS 47 | 2.55 |
| Vh PHE 56 | E GLN 271 | 3.873 |
| Vh SER 105 | E ASP 154 | 3.347 |
| **J9 Fv Light chain residues** | **E-dimer residues (E-E')** | **Distance ( all atom cutoff <5Å)** |
| Vl MET 98 | E' THR 68 | 3.223 |
| Vl GLN 27 | E' GLU 84 | 4.09 |
| Vl ASP 1 | E' THR 69 | 4.428 |
| Vl SER 93 | E' GLU 71 | 3.8 |
| Vl TRP 94 | E' LYS 246 | 2.764 |
| Vl PRO 95 | E' LYS 247 | 4.113 |
| Vl PRO 96 | E' LYS 247 | 4.14 |

| **J9 Fv Heavy chain residues** | **N153 glycan contact residues** | **Distance ( all atom cutoff <5Å)** |
|---|---|---|
| Vh SER 105 | E NAG 496 | 2.247 |
| Vh TYR 107 | E BMA 498 | 3.974 |
| Vh TYR 107 | E NAG 497 | 2.94 |
| Vh HIS 108 | E MAN 501 | 4.48 |
| Vh ASP 112 | E MAN 501 | 3.384 |
| **J9 Fv Light chain residues** | **N153 glycan contact residues** | **Distance ( all atom cutoff <5Å)** |
| Vl SER 56 | E MAN 501 | 3.146 |
| Vl TYR 49 | E BMA 498 | 3.545 |

| **J9 Fv Heavy chain residues** | **N67 glycan contact residues** | **Distance ( all atom cutoff <5Å)** |
|---|---|---|
| Vh ASN 64 | E’ NAG 497 | 3.9 |
| Vh ASP 63 | E' NAG 496 | 2.294 |
| **J9 Fv Light chain residues** | **N67 glycan contact residues** | **Distance ( all atom cutoff <5Å)** |
| Vl ILE 2 | E' NAG 496 | 2.939 |
| Vl VAL 3 | E' NAG 497 | 3.035 |
| Vl THR 5 | E' MAN 499 | 4.104 |

| **J9 Fv Light chain residues** | **Inter-raft contact residues** | **Distance ( all atom cutoff <5Å)** |
|---|---|---|
| Vl GLN 27 | E2' THR 303 | 1.5 |
| Vl SER 28 | E2' PRO 384 | 2.88 |
| Vl SER 26 | E2' ASP 329 | 4.4 |
| **J9 Fv Light chain residues** | **Inter-dimer contact residues** | **Distance ( all atom cutoff <5Å)** |
| Vl SER 28 | E2 THR 226 | 4.9 |

**Table S5.** Close contacts between D14.F25.S02 bnAb and DENV E-proteins based on high resolution cryo-EM maps (Cα-Cα distances < 9Å).

| F25 Fab | N153 glycan | Distance (Å) |
|---|---|---|
| **$V_H$ $C_A$** | | |
| Asp 114 | BMA 499 | 4.117 |
| Leu 115 | BMA 499 | 4.462 |
| Val 2 | NAG 497 | 4.545 |
| Gly 27 | NAG 497 | 6.008 |
| Gln 1 | NAG 497 | 6.590 |
| Leu 109 | NAG 496 | 6.911 |
| Ser 31 | NAG 496 | 7.191 |
| Tyr 32 | NAG 496 | 7.207 |
| Trp 116 | BMA 499 | 7.287 |
| Phe 113 | BMA 499 | 7.533 |
| Ile 111 | NAG 497 | 7.761 |
| Thr 28 | NAG 497 | 7.781 |
| **$V_L$ $C_A$** | | |
| Gly 59 | BMA 499 | 4.048 |
| Ser 58 | BMA 499 | 4.070 |
| Pro 57 | BMA 499 | 5.747 |
| Lys 47 | BMA 499 | 6.811 |
| Val 60 | BMA 499 | 6.483 |
| Leu 48 | BMA 499 | 7.686 |
| **F25 Fab** | **N67 glycan** | **Distance (Å)** |
| **$V_H$ $C_A$** | | |
| Gly 66 | NAG 499 | 2.948 |
| Arg 67 | NAG 499 | 4.537 |
| Ser 84 | NAG 499 | 6.392 |
| Gln 65 | NAG 499 | 6.517 |
| Val 68 | NAG 499 | 6.522 |
| Thr 69 | NAG 499 | 7.032 |
| **Intradimer Interactions** | | |
| **F25 Fab** | **E protein (E,E')** | **Distance (Å)** |
| **$V_H$ $C_A$** | | |
| Asp 108 | E' Gly 104 | 4.662 |
| Thr 28 | E Thr 155 | 4.897 |
| Thr 57 | E' Thr 70 | 5.512 |
| ALA 107 | E' CYS 105 | 5.675 |
| GLY 27 | E GLY 156 | 5.697 |

| | | |
|---|---|---|
| PHE 55 | E’ LYS 247 | 6.453 |
| SER 106 | E’ CYS 74 | 6.460 |
| SER 31 | E ASP 154 | 7.365 |
| **$V_L$ $C_A$** | | |
| SER 58 | E ASP 362 | 8.123 |
| **Inter-dimer interactions around 3 fold ($E_2$)** | | |
| **F25 Fab** | **E protein ($E_2$)** | **Distance (Å)** |
| **$V_L$ $C_A$** | | |
| TYR 97 | THR 226 | 5.625 |
| SER 96 | THR 226 | 5.830 |
| Tyr 97 | Gln 227 | 8.852 |
| **$V_H$ $C_A$** | | |
| ARG 10 | ALA 224 | 7.716 |
| **Inter-raft interactions around 5 fold (E’, E2’)** | | |
| **F25 Fab** | **E protein** | **Distance (Å)** |
| **$V_L$ $C_A$** | | |
| TYR 97 | E2’ PRO 384 | 2.741 |
| SER 96 | E2’ GLY 385 | 4.489 |
| GLY 31 | E’ GLU 174 | 4.995 |
| THR 98 | E2’ PRO 384 | 6.103 |
| GLY 30 | E’ GLU 174 | 6.211 |
| LYS 33 | E’ GLY 179 | 6.899 |
| TYR 32 | E’ GLY 179 | 7.333 |
| SER 69 | E’ THR 176 | 7.335 |

**Table S6.** Close contacts between amino acid side chains of D14.F05.S03 bnAb and DENV E-proteins based on high resolution cryo-EM maps.

| Fab ($V_H$ & $V_L$) residues | E protein (E, E', $E_2$) residues | Distances < 5Å |
|---|---|---|
| | **N67 glycan** | |
| ASP 60 ( $V_H$ ) | ASN 67 | 3.64 |
| | **N153 glycan** | |
| SER 104 ( $V_H$ ) | ASN 153 | 2.6 |
| | **Intra-dimer** | |
| GLN 26 ( $V_L$ ) | ASN 83 | 2.58 |
| GLN 26 ( $V_L$ ) | GLU 84 | 3.59 |
| SER 53 ( $V_H$ ) | LEU 277 | 4.2 |
| SER 53 ( $V_H$ ) | PHE 279 | 4.97 |
| ASP 60 ( $V_H$ ) | THR 66 | 4 |
| TYR 93 ( $V_L$ ) | THR 68 | 2.86 |
| TYR 93 ( $V_L$ ) | THR 69 | 4.78 |
| TYR 102 ( $V_H$ ) | TRP 101 | 4.95 |
| TYR 102 ( $V_H$ ) | GLY 104 | 4.02 |
| SER 104 ( $V_H$ ) | GLY 102 | 3.1 |
| SER 104 ( $V_H$ ) | GLY 152 | 4.94 |
| SER 104 ( $V_H$ ) | ASP 154 | 2.77 |
| SER 104 ( $V_H$ ) | THR 155 | 4.28 |
| SER 105 ( $V_H$ ) | ASN 103 | 3.39 |
| TYR 107 ( $V_H$ ) | ILE 113 | 3.96 |
| TYR 107 ( $V_H$ ) | THR 115 | 4.65 |
| TYR 107 ( $V_H$ ) | ASP 249 | 4.9 |
| TYR 107 ( $V_H$ ) | THR 70 | 3.8 |
| PHE 108 ( $V_H$ ) | LYS 246 | 4.27 |
| PHE 108 ( $V_H$ ) | SER 72 | 3.37 |
| PHE 108 ( $V_H$ ) | VAL 97 | 3.9 |
| PHE 108 ( $V_H$ ) | ASP 98 | 4.74 |
| PHE 108 ( $V_H$ ) | ARG 99 | 3.12 |
| | **Inter dimer** | |
| SER 27 | THR 226 | 2.5 |

**Table S7.** Contact surface area calculation using cryo-EM structures.

The contact area between Fab and the DENV E-dimer were calculated using the solvent accessible area (SA) of respective atomic models.

For calculating the contact area at Env-Fab interface:

[(Total SA of E-dimer + Total SA of Fab) – Total SA for complex] / 2

| DENV2 Virus-Antibody complex | Total Contact Area for both fabs (Å$^2$) | Contact Area due to Heavy Chains (Å$^2$) | Contact Area due to Light chains (Å$^2$) |
|---|---|---|---|
| J9 bnAb | 2496.5 | 1545 | 998.5 |
| D14.F25.S02 bnAb | 1890.5 | 1759.5 | 152.5 |
| D14.F05.S03 bnAb | 1959 | 1637.5 | 328.5 |

| DENV2 Virus-Antibody complex | Total SA of EM dimer of DENV (Å$^2$) | Total SA of Fabs (Å$^2$) | Total SA of EM dimer +Fab Complex (Å$^2$) | Total SA of Fab light chains (Å$^2$) | Total SA of Fab heavy chains (Å$^2$) | Total SA of EM dimer + Fab light Chain complex (Å$^2$) | Total SA of EM dimer+ Fab Heavy Chain complex (Å$^2$) |
|---|---|---|---|---|---|---|---|
| J9 bnAb | 59212 | 23194 | 77413 | 12712 | 14655 | 69927 | 70777 |
| D14.F25.S02 bnAb | 56932 | 24503 | 77684 | 12986 | 15201 | 69613 | 68614 |
| D14.F05.S03 bnAb | 53665 | 21483 | 71230 | 11734 | 13391 | 64742 | 63781 |
